# Protective anti-prion antibodies in human immunoglobulin repertoires

**DOI:** 10.1101/2020.02.05.933721

**Authors:** Assunta Senatore, Karl Frontzek, Marc Emmenegger, Andra Chincisan, Marco Losa, Regina Reimann, Geraldine Horny, Jingjing Guo, Sylvie Fels, Silvia Sorce, Caihong Zhu, Nathalie George, Stefan Ewert, Thomas Pietzonka, Simone Hornemann, Adriano Aguzzi

**Author notes:** Correspondence to: Dr. Adriano Aguzzi or Dr. Simone Hornemann, Institute of Neuropathology, University Hospital Zurich, Schmelzbergstrasse 12, CH-8091 Zürich, Switzerland. equal contribution (Karl Frontzek, Marc Emmenegger, Simone Hornemann).

## Abstract

Prion immunotherapy may hold great potential, but antibodies against certain PrP epitopes can be neurotoxic. Here we identified >6000 PrP-binding antibodies in a synthetic human Fab phage display library, 49 of which we characterized in detail. Antibodies directed against the flexible tail of PrP conferred neuroprotection against infectious prions. We then mined published repertoires of circulating B cells from healthy humans and found antibodies similar to the protective phage-derived antibodies. When expressed recombinantly, these antibodies exhibited anti-PrP reactivity. Furthermore, we surveyed 48’718 samples from 37’894 hospital patients for the presence of anti-PrP IgGs, and found 21 high-titer individuals. The clinical files of these individuals did not reveal any enrichment of specific pathologies, suggesting that anti-PrP autoimmunity is innocuous. The existence of protective anti-prion antibodies in unbiased human immunological repertoires, combined with the reported lack of such antibodies in carriers of disease-associated *PRNP* mutations, suggests a link to the low incidence of spontaneous prion diseases in human populations.

## Introduction

Many neurodegenerative syndromes, including prion diseases, Alzheimer’s disease and Parkinson’s disease, go along with the accumulation of misfolded and aggregated proteins in the central nervous system. Antibodies against such proteins may be beneficial^1^, e.g. by opsonizing pathological aggregates and mediating their degradation by phagocytic cells^2, 3^. While the clinical effectiveness of antibody-based therapies against neurodegenerative diseases is still being debated^4^, there is ample evidence that both active immunization and passive antibody transfer can effectively clear pathological aggregates in preclinical animal models and, to some extent, in affected humans.

According to the protein-only hypothesis, the prion is an infectious particle consisting of PrP^Sc^, an aggregated and proteinase K (PK) resistant isoform, of the cellular prion protein PrP^C 5^. PrP^C^ consists of a C-terminal globular domain (GD) and an N-terminal flexible tail (FT) which includes the octapeptide repeat (OR) region, two cationic charge clusters (CC1 and CC2) and a hydrophobic core (HC)^6^. The CC1 domain of PrP^C^ participates in Schwann cell maintenance by activating the G protein-coupled receptor Adgrg6^7, 8^. While PrP^C^ deficient mice are only mildly affected, PrP^Sc^ necessitates PrP^C^ for its propagation^9^ and prion toxicity is transduced by PrP^C^ onto target cells^10, 11^. Therefore, suppression of PrP^C^ by means of anti-PrP^C^ antibodies represents a rational strategy against prion diseases.

Here we panned a synthetic human antibody phage display library to explore the presence of PrP-binding antibody fragments (Fabs)^12^. To identify rare antibodies to poorly antigenic epitopes that may be overlooked by conventional screening technologies, we performed “next-generation” sequencing (NGS) of panning outputs after phages selections^13^. Several anti-PrP binders were identified and found to antagonize prion toxicity. What is more, mining of published human antibody repertoires identified sequences similar to an anti-prion phage-derived Fab which, when expressed, acted as functional PrP binders. Lastly, the interrogation of a large unselected hospital cohort (n = 37’894) highlighted individuals with high-titer anti-PrP autoreactivity whose clinical presentation was heterogeneous, yet unrelated to known features of prion diseases. Therefore, anti-prion immunity can exist in human communities and is seemingly innocuous.

## Results

### Phage display selection strategy for anti-PrP Fabs

We used three rounds of phage display to screen two synthetic human Fab phage display libraries (Extended data Fig. 1a) with short (8-10 aa) and long (12-20 aa) heavy-chain complementarity-determining regions 3 (HCDR3). These libraries were constructed to mimic human antibody repertoires by combining frameworks from human germline sequences with diversified HCDR3 whose design approximated the natural gene sequences in human repertoires as compiled in the IMGT database^14^. The first and second biopanning rounds were performed against full-length recombinant mouse PrP (recPrP_23-231_) to enrich for Fabs covering a large variety of PrP epitopes. To further select Fabs recognizing specific PrP epitopes, a third panning round was conducted against several different antigens in parallel, including recPrP_23-231,_ recPrP_23-110_ spanning the FT, the GD (recPrP_90-231_ and recPrP_121-231_) and synthetic peptides representing CC1_23-50_ spanning the CC1, N-OR_39-66_ and F-OR_51-91,_ containing the OR, and CC2-HC_92-120_, spanning CC2 and the HC.

**Figure 1:**
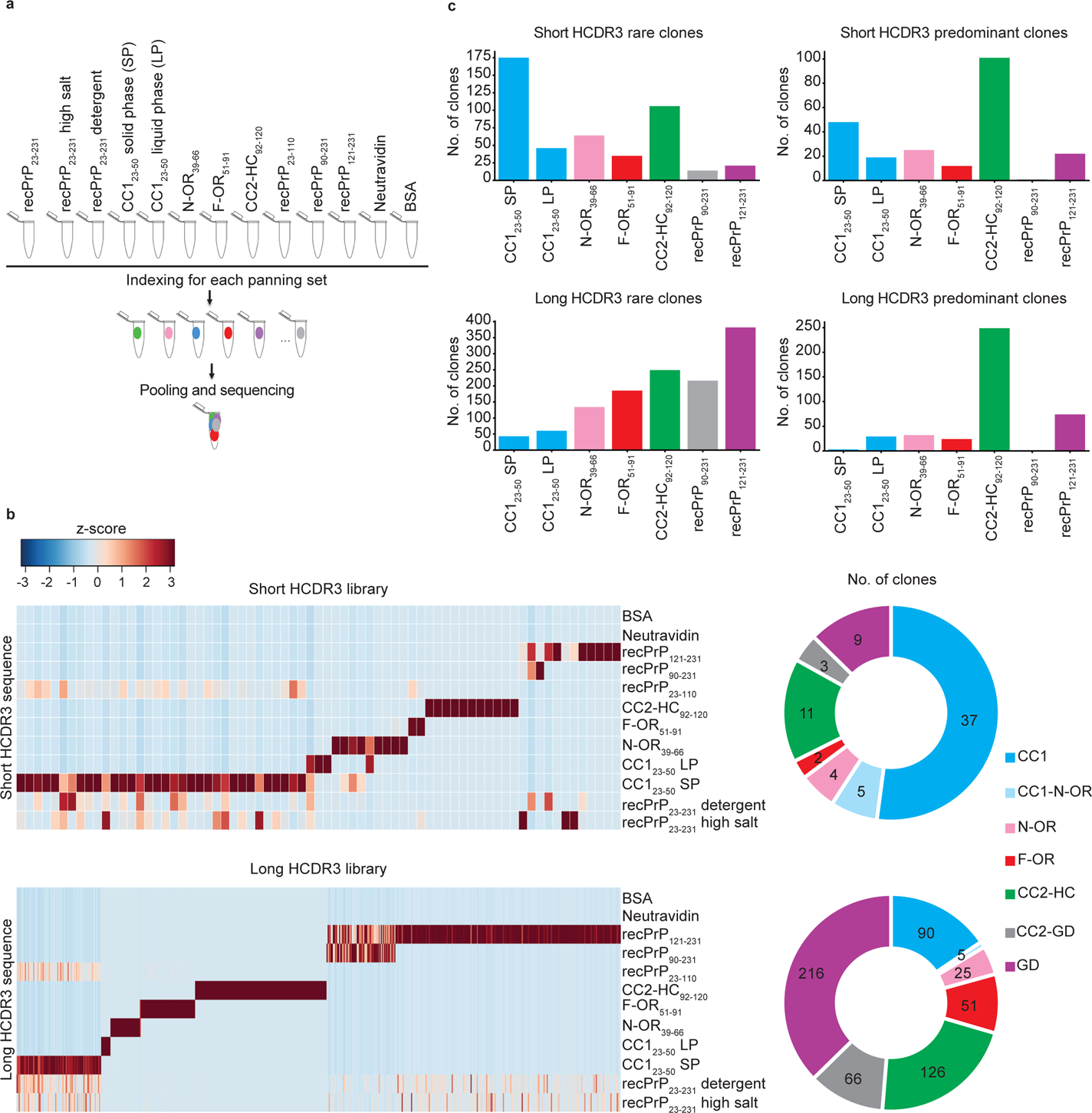
Identification of PrP binders. (a) Preparation of HCDR3-Fab DNA libraries for NGS analysis. Three rounds of panning were performed against several mouse recPrP fragments, synthetic PrP peptides, recPrP_23-231_, and recPrP_23-231_ using various washing conditions and against CC1_23-50_ in solid and liquid phase. DNA sequences of selected phages from the panning output were indexed by sequential PCR reactions, pooled and analyzed by NGS for their HCDR3 sequences. Control: Panning on Neutravidin and BSA. **(b)** Heatmap representation of enriched HCDR3 sequences across the different panning sets for the short-HCDR3 and long-HCDR3 (upper and lower panel, respectively) phagemid libraries. HCDR3 sequences were selected based on NGS counts in 100’000 analyzed sequences (Z-score values; Table S2) and clustered according to the NGS-binding profiles. Red and blue: high and low number of NGS counts of the HCDR3 sequence, respectively. Donut charts (the right side) of each heat map indicate the number of clones with NGS-identified HCDR3 for a predicted PrP epitope. **(c)** Bar graphs showing the number of rare (one count in mouse recPrP_23-231_ panning) and predominant (count ≧ 20) clones binding to the various PrP regions.

Only few existing antibodies bind to the natively unstructured CC1_23-50_^15^. To optimize our chances to identify CC1_23-50_ binders, and to avoid misfolding artefacts caused by nonspecific plate adsorption, the selection against the biotinylated CC1_23-50_ peptide was conducted in solid phase and by liquid phase panning followed by neutravidin (Neu)-mediated capture. Liquid-phase panning was also performed for other biotinylated peptides (N-OR_39-66_, F-OR_51-91_ and CC2-HC_92-120_). Furthermore, we performed panning rounds using a matrix of stringent washing conditions (Extended data Table 1) to also include an affinity read-out to the analysis of the NGS screening.

**Table 1:** Dissociation rate (koff) and equilibrium (KD) constants of the PrP binding Fabs determined by SPR

|  | **k<sub>off</sub> (1/s)** | **K<sub>D</sub> (M)** |
| --- | --- | --- |
| **Fab_POM1** | 2.96E-04 | 2.01E-09 |
| **Fab1** | 2.88E-03 | 1.21E-08 |
| **Fab2** | 1.58E-03 | 1.01E-08 |
| **Fab3** | 2.46E-03 | 7.72E-09 |
| **Fab4** | 1.93E-03 | 1.41E-08 |
| **Fab6** | 1.44E-03 | 9.89E-09 |
| **Fab7** | 2.41E-03 | 1.76E-08 |
| **Fab8** | 3.62E-03 | 1.79E-08 |
| **Fab10** | 2.61E-03 | 1.41E-08 |
| **Fab12** | 3.20E-03 | 1.71E-08 |
| **Fab13** | 4.98E-03 | 4.56E-08 |
| **Fab15** | 3.76E-03 | 1.27E-08 |
| **Fab25** | 2.42E-03 | 4.71E-08 |
| **Fab28** | 1.08E-03 | 1.28E-08 |
| **Fab29** | 3.15E-03 | 1.04E-08 |
| **Fab30** | 1.76E-03 | 4.76E-09 |
| **Fab32** | 2.90E-03 | 1.50E-08 |
| **Fab35** | 2.29E-03 | 1.75E-08 |
| **Fab41** | 2.65E-03 | 1.95E-08 |
| **Fab44** | 3.18E-03 | 1.21E-08 |
| **Fab46** | 3.39E-03 | 1.46E-08 |
| **Fab48** | 2.45E-03 | 1.06E-08 |
| **Fab52** | 4.12E-03 | 2.98E-08 |
| **Fab53** | 1.73E-03 | 9.79E-09 |
| **Fab61** | 3.97E-03 | 2.06E-08 |
| **Fab69** | 1.36E-02 | 9.50E-08 |
| **Fab71** | 2.45E-03 | 6.20E-09 |
| **Fab72** | 2.14E-03 | 7.14E-09 |
| **Fab74** | 1.57E-02 | 4.85E-08 |
| **Fab75** | 3.67E-03 | 2.64E-08 |

### Next Generation Sequencing (NGS) of anti-PrP Fabs

The HCDR3 domains can contribute crucially to antigen binding^16^. Sequencing of the outputs of the third panning rounds yielded 4,847 and 11,948 unique HCDR3 sequences in 100’000 analyzed sequences for the short and long HCDR3 Fab libraries, respectively. We excluded all HCDR3 that had any counts in the negative-control outputs (Neu and BSA panning) and retained only clones with ≥ 1 read in the recPrP_23-231_ output. These constraints reduced the unique HCDR3 sequences to 1,173 and 4,832 anti-PrP Fabs, respectively. We then compared the read counts of each HCDR3 between panning to recPrP_23-231_ and the different PrP domains. For each HCDR3, we considered the enrichment of NGS counts in a panning output as reflecting the binding to the respective PrP peptide/fragment used in the panning. As an example of the stringent sorting criteria for epitope binding profile determination (Extended data Table 2), all HCDR3 having counts > 0 in CC1_23-50_ and in recPrP_23-110_ outputs, and count = 0 in all other panning outputs, were classified as specific PrP binders in the CC1_23-50_ region. HCDR3 sequences were clustered based on their NGS-binding profile and found to represent a highly diverse collection of anti-PrP Fabs (Fig. 1a, b). Predominant clones binding to the high antigenic epitopes, defined as those with ≥ 20 NGS counts in recPrP_23-231_ panning, were mostly directed against the CC2-HC_92-120_ in both HCDR3 libraries (Fig. 1c). Rare clones against less antigenic epitopes, i.e. displaying only one count in recPrP_23-231_ panning, were predominantly showing an NGS-binding profile to the CC1_23-50_ and the GD domains (Fig. 1c).

We retrieved clones of interest, as identified by the HCDR3 read profile in NGS, by overlapping PCR from the third-round polyclonal phagemid DNA (Extended data Fig. 1b-e). In one instance we designed primers to an HCDR3 sequence with an NGS-binding profile to the CC2-HC_92-120_ epitope (NGS read enrichment in the CC2-HC_92-120_ panning as compared to reads in panning to other PrP domains, Extended data Fig. 1b) and retrieved entire Fab sequences by PCR (Extended data Fig. 1c). The retrieved Fab, designated as FabRTV, was cloned into an E. coli expression vector, Sanger-sequenced and purified by immobilized metal ion affinity chromatography (IMAC) (Extended data Fig. S1d). Enzyme-linked immunosorbent assay (ELISA) confirmed the NGS-binding profile of FabRTV to the CC2-HC_92-120_ domain (Extended data Figs. 1b, e).

### Identification of anti-PrP Fabs by ELISA screening

As a complementary approach, we screened 4416 clones (randomly selected by an automated colony picker) by ELISA. We then selected 312 hits reactive to recPrP_23-231_, to recPrP fragments, or to synthetic biotinylated peptides (>10 or >5-fold over background, while displaying a Neu signal <5-fold over background). From those, eighty confirmed anti-PrP Fabs were Sanger-sequenced, produced in E. coli and epitope-mapped by ELISA (Extended data Fig. 2). Out of 49 Fabs, 4 targeted CC1_23-50_, 15 OR_51-91_, 22 CC2-HC_92-120_ and 8 the GD (Extended data Table 3). The abundance of CC2-HC_92-120_ targeting Fabs is in agreement with the NGS analysis showing that CC2-HC_92120_ binders are the predominant clones. For 35 of these Fabs, the ELISA binding results confirmed the epitope binding profile determined by NGS analysis.

**Figure 2:**
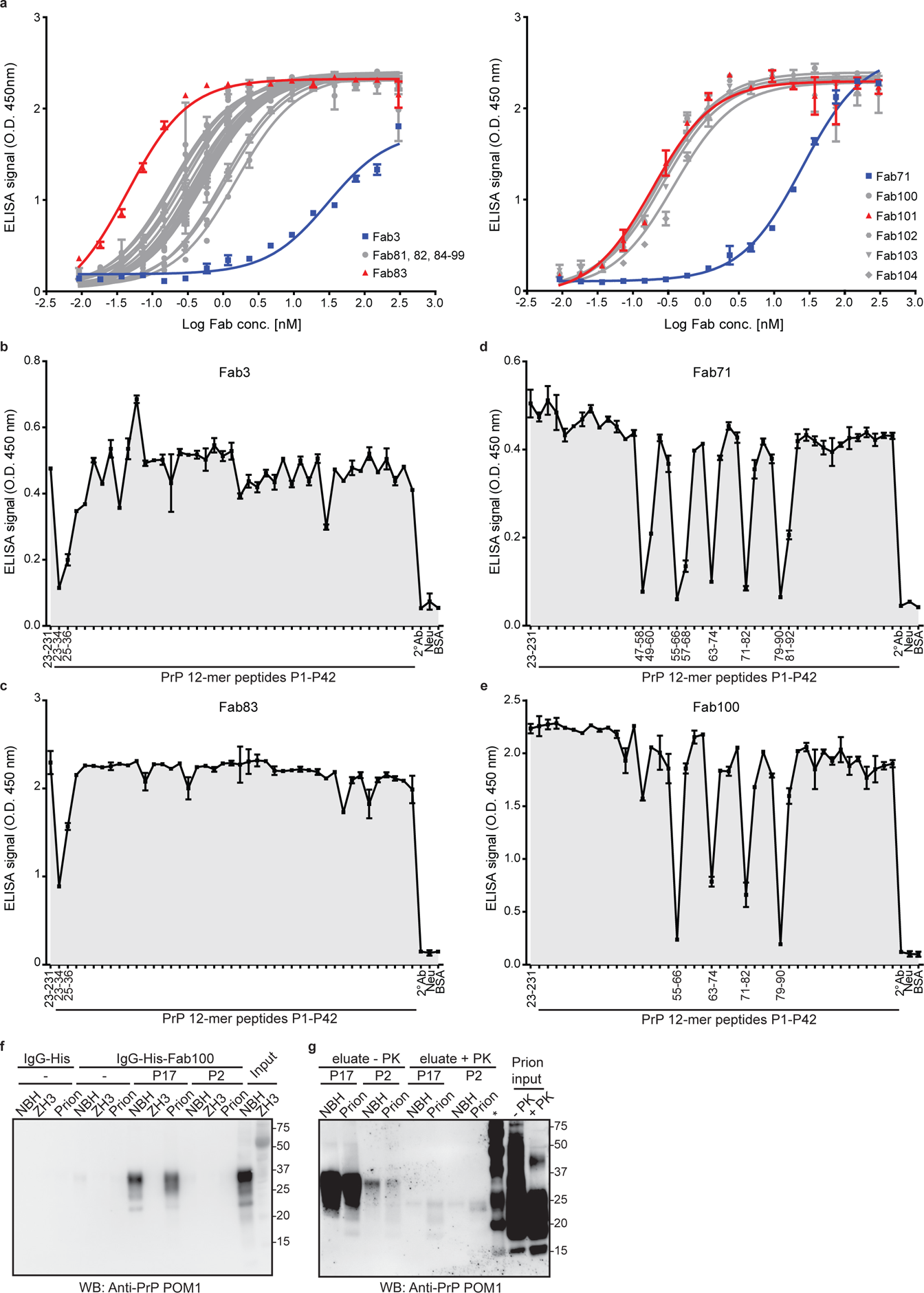
Binding specificities of retrieved anti-PrP Fabs by ELISA and immunoprecipitation. (a) ELISA titration curves (OD at 450nm) of Fab3 (red; left panel) and Fab71 (red; right panel) against mouse recPrP_23-231_ compared to their best affinity-matured versions, Fab83 and Fab101 (blue; EC_50_ values in Table S3). ELISA data here and henceforth were performed in duplicates. Data presented in dot plots represent the mean ± sem (here and henceforth). Curves were fitted by non-linear least-squares regression analysis for statistical analysis: log(agonist) vs. response. **(b-e)** FT-peptide competition ELISA to map the epitopes of the indicated Fabs. Peptides that strongly inhibit the binding of the Fabs to recPrP_23-231_ are indicated by their residue numbers in the PrP sequence and reflect the respective binding epitopes. Positive control: recPrP_23-231;_ negative controls: Neu, BSA, and the secondary antibody (2° Ab). **(f)** IgG-His-Fab100 coupled beads efficiently immunoprecipitated wtPrP^C^ from non-infectious brain homogenate (NBH) and total wtPrP from brains of prion-infected mice (prion), but not from brains of *Prnp*^ZH3/ZH3^ (ZH3) mice. WtPrP was eluted by competition with the epitope-targeting peptide P17, but not with the unrelated peptide P2. Eluates were visualized by western blotting. Control: IgG-His. Molecular sizes are presented in kDa. **(g)** Same as (f) for NBH and prion to confirm the specific immunoprecipitation and detection of PrP^Sc^ with the peptide P17 from brains of prion-infected mice. +PK: digested with PK; -PK: non-digested with PK.

### Kinetic measurements by surface plasmon resonance (SPR) and affinity maturation

The K_D_s (determined by SPR as k_off_/k_on_ ratio) of all tested Fabs to recPrP_23-231_ were in the range of 10-100 nM (Table 1) with fast dissociation rate constants (k_off_ > 10^-3^ [s^-1^]).

To optimize the binding properties of Fab3 and Fab71, to CC1_23-50_ and OR_51-91_ respectively, we used affinity-maturation libraries in which the parental HCDR2 and LCDR3 loops were replaced with pre-built highly diversified cassettes. In addition, phage display selections were repeated with more stringent conditions than in the original selection. To ensure retention of specificity to the respective epitopes, panning against recPrP_23-231_ was alternated with panning to the CC1_23-50_ fragment for Fab3 and to the OR_51-91_ fragment for Fab71. Again, selected Fabs were screened by NGS and ELISA. After high-throughput off-rate and on-rate ELISA screening, 19 and 5 affinity-matured versions of Fab3 (Fab81-Fab99) and Fab71 (Fab100-Fab104) were identified, respectively. These clones were also prevalent in the NGS dataset. With this strategy, the EC_50_s of the affinity-matured Fabs were improved by 2-3 logs over the parental Fabs (Fig. 2a and Extended data Table 4).

#### Epitope confirmation and mapping

Next, we confirmed the binding behavior of the different Fabs to biotinylated FT-PrP peptides CC1_23-50_, N-OR_39-66_, F-OR_51-91_ and CC2-HC_92-1_20 by SPR (Extended data Fig. 3). For all tested Fabs, the specificity of the targeted epitope matched the ELISA epitope profiling. Being unstructured, the FT displays many linear epitopes^17^. Therefore, we additionally mapped the epitopes of the anti-FT Fabs by competition ELISA using overlapping dodecameric PrP peptides (each shifted by 2 residues) spanning residues 23-120. The binding of Fab3 and of its affinity-matured version Fab83 to immobilized recPrP_23-231_ was blocked by peptides P1 and P2 (residues 23-34 and 25-36, respectively; Fig. 2b,c), pointing to the KKRPKPG polybasic stretch as their minimal epitope.

**Figure 3:**
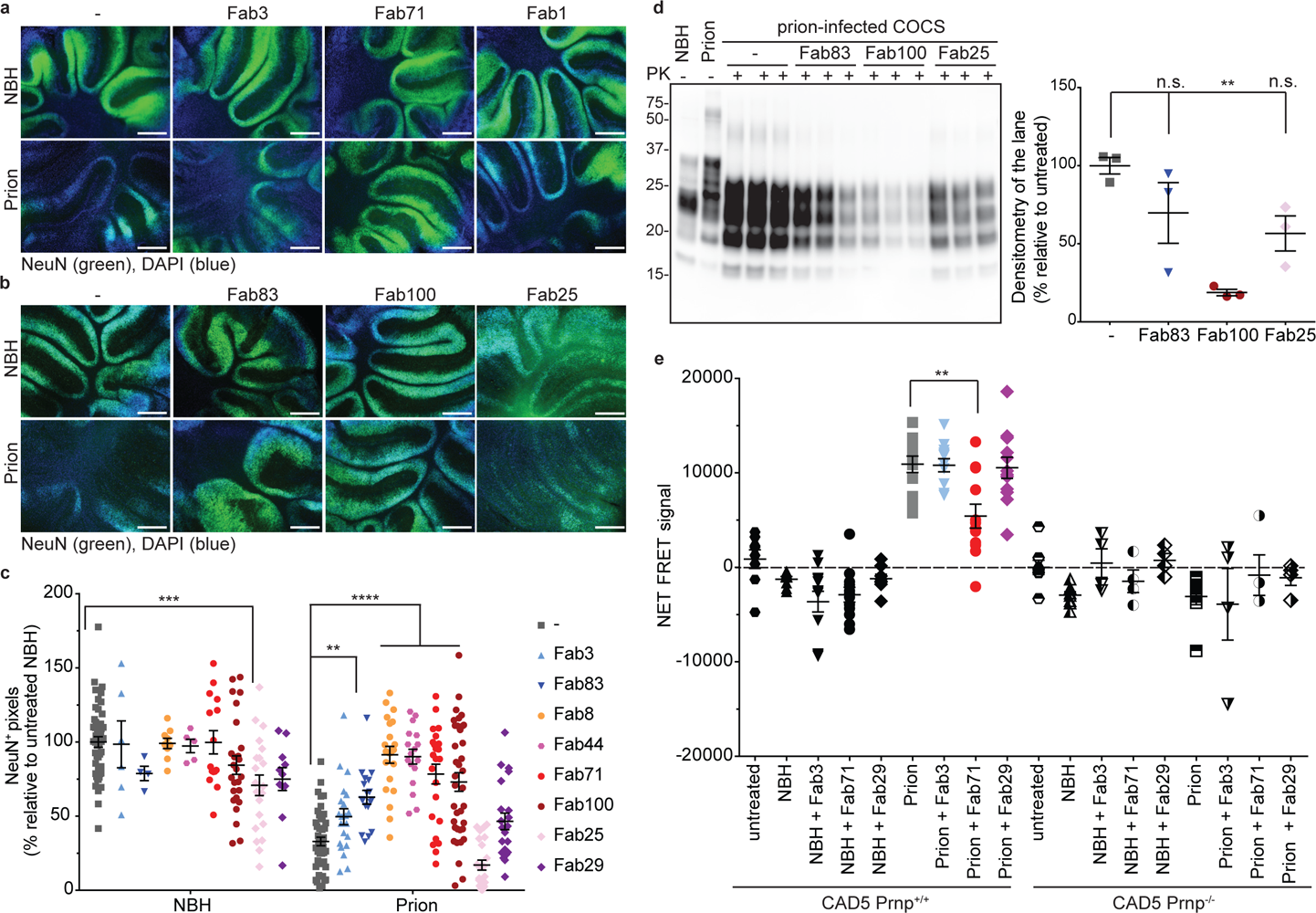
Fabs targeting the CC1 and OR of PrP^C^ prevent prion-induced neurotoxicity in COCS. (a) Fluorescence micrographs of *tg*a*20* COCS chronically exposed to prions and cultured in the presence of either Fab3 (CC1_23-50_), Fab71 (OR_51-91_) or Fab1 (GD) for 45 days. Scale bar: 500 μm. **(b)** Same as (a), but in the presence of the affinity-maturated antibodies. **(c)** NeuN immunofluorescence coverage of prion-infected COCS cultured in the presence of the Fabs. Antibodies targeting the CC1_23-50_ and the OR_51-91_ afforded protection against neurodegeneration. Each dot represents a cerebellar slice for all treatment groups. **(d)** Western blot analysis of COCS lysates. Prion-infected COCS treated with Fab100, but not with Fab83 and Fab25 showed reduced levels of PrP^Sc^. n = 3 biological replicates. **(e)** TR-FRET for the detection of PrP^Sc^ in prion-infected CAD5 *Prnp^+/+^* cells after treatment with selected anti-PrP Fabs. Treatment of the cells with Fab71 (OR_51-91_), but not with Fab3 (CC1_23-50_) or Fab29 (GD), reduced PrP^Sc^ levels compared to untreated, prion-infected cells. The FRET signal of untreated CAD5 *Prnp^-/-^* cells was set to zero. Negative controls: NBH and CAD5 *Prnp^-/-^* cells. n = 4-12. One way (d) and two-way ANOVA (c), respectively, followed by Bonferroni’s post-hoc test were used for statistical analysis; ** p<0.01; *** p<0.001; **** p<0.0001; n.s.: not significant.

The anti-OR_51-91_ Fab8, Fab12, Fab71 and the affinity maturated Fab100 recognized the sequence, WGQPHGGG(S)WGQ, which is repeated 4 times in PrP (residues 55-66, 64-74, 72-82 and 80-90). Additionally, Fab8, Fab12, and Fab71 also targeted NRYPPQGGTWGQ at the very beginning of the OR domain (residues 47-58; Fig. 2d,e and Extended data Fig. S4). Fab44 recognized the sequence WGQPHGG within the OR (Extended data Fig. 4). For Fab7, Fab10, Fab41 and Fab52, we could not identify any PrP peptide that abrogated the ELISA signal to recPrP_23-231_. This may point to the presence of conformational/discontinuous epitopes. The CC2-HC_92-120_ directed Fab13, Fab53, Fab61 and Fab69 all recognized residues 93-100 (GTHNQWNK; Extended data Fig. 4).

**Figure 4:**
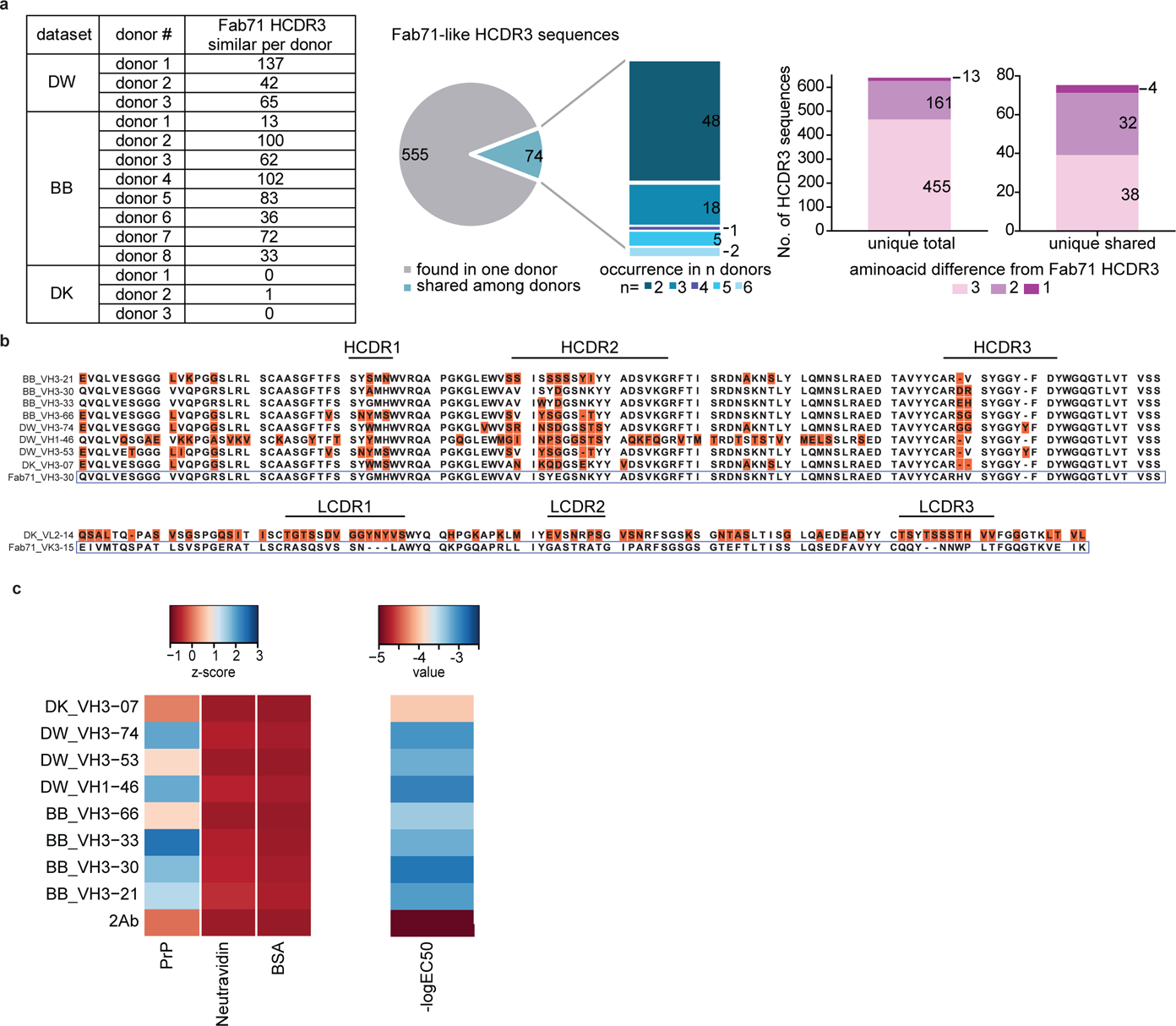
Identification of anti-PrP Fab71 analogous antibodies in human antibody repertoires. (a) Fab71-similar HCDR3 sequences in three different NGS datasets (DW, BB, and DK) of human antibody repertoires from healthy donors. Pie chart and stacked bar plot (middle) indicate the occurrence of the identified Fab71 similar HCDR3s: 555 out of 629 identified sequences were only found in one donor, while 74 sequences were shared, occurring in 2 up to 6 donors. Stacked bar plots (right) report the number of sequences differing to Fab71 HCDR3 by 3, 2 or 1 residues. In total, 13 sequences differed by only one residue. Among them, 4 sequences were shared between different donors. **(b)** Sequence alignment of Fab71 VH3-30 with HCDR3 loops that differ from Fab71 by ≤ 3 residues (upper panel). Lower panel: sequence alignment of Fab71 light chain VK3-15 with DK_VL2-14 that is naturally paired with DK_VH3-07. **(c)** Left: Heat map showing the binding specificity (Z-scores) of selected Fab71 human analogous antibodies to human recPrP_23-230_ compared to the negative controls (BSA and Neu). Right: Heat map representing the reactivity (-logEC50) obtained from dose-dependent ELISA binding curves of the analogous antibodies to human recPrP_23-230_.

In addition, Fab83, Fab100, Fab53 and Fab74 were also able to detect wild-type PrP^C^ (wtPrP^C^) in brain homogenates (BH) from wild-type (C57BL6/J) and *tg*a20 mice^18^ overexpressing wtPrP^C^, but not in the brains of mice expressing PrP deletion mutants lacking the respective epitopes (Extended data Fig. 5b).

**Figure 5:**
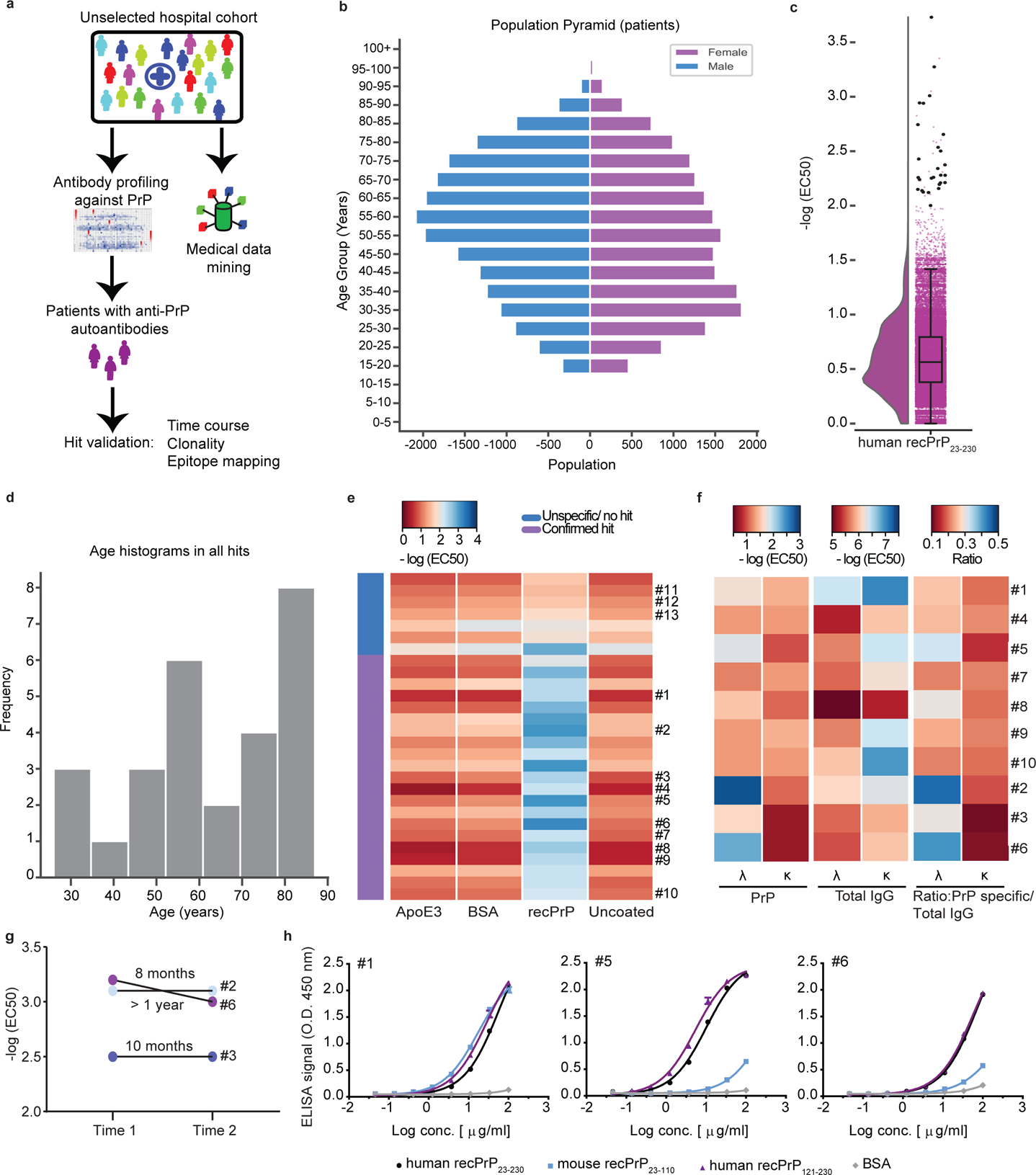
Identification of anti-PrP autoantibodies in an unselected hospital patient cohort. (a) Schematic workflow for the HTS of patient plasma samples for anti-PrP autoantibodies. **(b)** Age pyramid, separated for females and males, of the 37,894 unselected hospital patients tested in the HTS. **(c)** Boxplot with half-violin plot displaying the distribution of autoantibody reactivity in 48,718 plasma samples. Twenty-seven samples over the reactivity threshold of -log(EC50) ≥ 2 and fitting error < 20% were considered as hits in the primary screen (black dots: single samples). **(d)** Frequency distribution of hits based on age. **(e)** Heat map representation for the validation of the hits from the primary screen (21 out of 27 hits). Confirmed hits displayed selective reactivity against human recPrP_23-230_. Controls: Patients #11-13, ApoE4 and BSA. Red: high reactivity; blue: low reactivity. **(f)** Heat maps representing autoantibody clonality of hits analyzed by ELISA. Autoantibodies were mostly constituted by lambda light chains (λ), while the kappa light chains (κ) showed prevalence in the total IgG fraction. **(g)** Determination of antibody reactivity (-log(EC50)) for three individuals at two different time points. Individuals were selected by displaying strong anti-PrP IgG reactivity in the first test (Time 1). Reactivity was maintained for even more than one year (Time 2). **(h)** Purified IgGs from patients with strong anti-PrP reactivity were tested against PrP variants and BSA (negative control) in dose-dependent ELISAs. Patient #5 and #6 showed binding mostly to the human recPrP_121-230_. Patient #1 displayed a polyreactive pattern with binding to the human recPrP_121-230_ and mouse recPrP_23-110_.

The CC1_23-50_ binder Fab83 did not detect PrPΔ25-50^19^ and PrPΔ32-93^20^ on Western blots, nor did it detect cell expressed PrPΔ23-27, PrPΔ23-31 and PrP mutated in the polybasic stretch from KKRPK to KPRKK by ELISA (Extended data Fig. 5c). Lysine-to-alanine substitutions within the KKRPK sequences reduced the binding of Fab83. Hence the Fab83 epitope comprises both lysine K24 and proline P26 in the N-terminal KKRPK stretch (PrP23-27) and additional residues in the PrP32-36 segment. Alternatively, KKRPK could have become buried in the truncated version PrPΔ32-93.

Consistent with the peptide epitope mapping, Fab100 did not react with the OR-deleted PrPΔ32-90 and PrPΔ51-90^19^. In addition, Fab53 did not detect PrPΔ94-110^7^, whereas the anti-GD Fab74 recognized all FT-modified versions of PrP. Most of the tested Fabs also cross reacted with human recPrP_23-230_ (Extended data Fig. 5a).

We then performed immunoprecipitations of wtPrP^C^ from BH of wt mice and from mice expressing different PrP deletion mutants, with Fab83 (CC1_23-50_) and Fab71 (OR_51-91_) (Extended data Fig. 5d,e). Fab83 and Fab71 immunoprecipitated wtPrP^C^ but not PrPΔ25-50 and PrPΔOR with deletion of the Fab binding sites as predicted from the ELISA. wtPrP^C^ was eluted by the P1 peptide containing the Fab83 binding site within the CC1_23-50_ but not by the P15 peptide within the OR_51-91_ (Extended data Fig. 5d). Similarly, Fab71 immunoprecipitated wtPrP^C^ was eluted from the Fab71-beads complex by the epitope-mimicking peptide P17 in the OR_51-91_, but not by the P3 peptide within the CC1_23-50_ (Extended data Fig. 5e).

We then tested Fab100 for immunoprecipitation of PrP^Sc^ from prion infected brain homogenates (Fig. 2f). Fab100 immunoprecipitated PrP^C^ and PrP^Sc^ from NBH and prion-infected brains, respectively. Elution under native conditions was achieved using peptide P17 (within OR_51-91_) but not P2 (within CC1_23-50_), confirming the specificity of Fab100 for the OR of both PrP^C^ and PrP^Sc^. Proteinase-K (PK) digestion assays confirmed the presence of PrP^Sc^ in the eluted fractions from prion-infected BH after immunoprecipitation by Fab100 (Fig. 2g).

#### Validation by flow cytometry and immunostaining

We also assessed the binding of a panel of Fabs to cell-surface wtPrP^C^ by flow cytometry using the murine neuroblastoma cell line CAD5. For each Fab, we compared the mean fluorescence intensity (MFI) between CAD5 *Prnp*^+/+^ and *Prnp*^-/-^ cells (Extended data Fig. 6a-b). All CC1_23-50_ binders, the majority of OR_51-91_ binders and CC2-HC_92-120_ binders discriminated between *Prnp^+/+^* and *Prnp*^-/-^ CAD5 cells, with Fab71 being the best performer. Of the GD binders, only Fab74 showed a differential signal, suggesting that the GD of membrane-bound wtPrP^C^ is less accessible to antibodies than the FT. Binders were tested by immunofluorescence on paraffin-embedded brain sections of C57BL6/J, *Prnp*^ZH3/ZH3^ and *tg*a*20* mice. Fab3, Fab71 and Fab74 detected murine wtPrP^C^ (Extended data Fig. 6c) with high specificity.

#### Activity of Fabs in models of prion disease

Antibodies that bind to the FT have been demonstrated to be neuroprotective, whereas those directed against certain epitopes of the GD are invariably toxic^11, 21^. We therefore asked whether Fabs against CC1_23-50_ (Fab3 and Fab83), the OR_51-91_ (Fab8, Fab44, Fab71 and Fab100) and the GD (Fab25, Fab1 and Fab29) may counteract neurotoxicity in prion-infected cerebellar slices (COCS)^11, 22^. *Tg*a20 COCS were exposed to Rocky Mountain Laboratory prions (passage #6, RML6) or to NBH and cultured in the presence or absence of the different Fabs (550 nM). At 45 days post-infection (dpi), prion-infected COCS showed conspicuous neurodegeneration (Fig. 3a-c). The GD binder Fab25 showed intrinsic neurotoxicity as indicated by loss of NeuN immunoreactivity in NBH-treated COCS. All tested OR_51-91_ binders, but none of the GD binders prevented prion-induced neurotoxicity. Fab3 (binding to CC1_23-50_) did not prevent prion neurotoxicity, whereas its affinity matured derivative Fab83 was effective (Figure 3a-c).

We then investigated the effects of the Fabs on prion replication and PrP^Sc^ accumulation (Fig. 3d). Treatment with the anti-OR_51-91_ Fab100, but not by anti-CC1_23-50_ Fab83, reduced PrP^Sc^ in prion-infected COCS, although both were neuroprotective. In slices treated with the GD Fab25, PrP^Sc^ levels were not significantly different from untreated prion-inoculated slices.

We also assessed the ability of the Fabs to arrest prion replication and/or PrP^Sc^ accumulation in prion susceptible cells. CAD5 *Prnp^+/+^* and CAD5 *Prnp^-/-^* cells were exposed to prions or NBH and treated with 10 µg/ml of either CC1_23-50_ binder Fab3, OR_51-91_ binder Fab71 or the GD targeting Fab29. Fabs were added to the medium 1 hour after infection and after every splitting. At 14 dpi, PrP^Sc^ content was determined using a homogenous-phase PrP^Sc^ time resolved (TR) FRET quantification immunoassay after proteolytic PrP^C^ removal. PrP^Sc^ was seen by TR-FRET in prion infected CAD5 *Prnp^+/+^* but not in CAD5 *Prnp^-/-^* cells indicating that the initial prion inoculum had been diluted below detectability. Fab83 did not reduce PrP^Sc^ level in CAD5 *Prnp^+/+^* cells despite being protective in COCS (Extended data Fig. 7). Fab71 and Fab100, but not Fab3 and Fab29, substantially lowered PrP^Sc^ in prion infected CAD5 *Prnp^+/+^* cells (Fig. 3e and Extended data Fig. 7). Hence OR_51-91_ binders exerted their neuroprotective activity by reducing prion accumulation.

#### Identification of anti-PrP antibodies from human antibody repertoires similar to the phage-derived Fabs

The Fabs were derived from a synthetic library mimicking the human antibody repertoire, raising the question whether analogous antibodies might be present in bona fide human repertoires. The heavy and light-chain frameworks of Fab71 correspond to human germlines VH3-30 and Vκ3-15, respectively, whereas its HCDR3 originates from the library randomization strategy. We therefore searched for human antibodies harboring HCDR3 amino acid sequences similar to that of Fab71. We examined large-scale repertoire datasets with billions of sequences that were generated recently by sequencing heavy-chain (VH; here referred to as DW^23^ and BB^24^ datasets) and natively paired heavy/light-chain variable regions (VH:VL; DK dataset)^25^ from circulating naïve and memory B cells of healthy donors. While no exact match to Fab71 HCDR3 was found, twelve out of fourteen analyzed donors displayed 629 HCDR3 sequences that differed from Fab71 HCDR3 by ≤ 3 residues (Fig. 4a). Among them, 74 HCDR3 sequences occurred in ≥ 2 subjects, four of which differed from Fab71 HCDR3 by only one residue (Fig. 4a).

We selected eight heavy chain variable regions similar to Fab71. In one of these, HCDR3 deviated from Fab71 in just two residues and matched the Fab71 VH3-30 segment. Three further HCDR3 differed from Fab71 HCDR3 by only one residue and two of those HCDR3 appeared in antibodies using different V genes (VH3-21 and VH1-46) from three subjects (Fig. 4b). We expressed these heavy chains variable regions in bacteria along with the Fab71 light chain to purify Fabs and test their reactivity to human recPrP_23-230_. In the case of DK_VH3-07, where VL sequence information was available from the VH:VL paired sequencing dataset, we used VL_2-14. All selected naturally occurring Fab71-like antibodies, except the DK_VH3-07, recognized human recPrP_23-230_ by ELISA specifically (Fig. 4c). BB_VH3-30, DW_VH1-46 and DW_VH3-74 Fabs, displayed the highest apparent affinities (EC50 = 1.21 µM, EC50 = 1.07µM and EC50 = 0.98 µM respectively; for DK_VH3-07 Fab, which did not bind to human recPrP_23-230_, calculated EC50 = 62.8 µM). While these affinities are low, they may be greatly raised by dimeric cooperativity in bivalent antibodies.

#### Identification of natural anti-PrP autoantibodies in an unselected hospital cohort

To assess the validity of the above findings, we interrogated a large cohort of human individuals for naturally occurring antibodies against PrP (Fig. 5a and Extended data Fig. 8). We performed an automated microELISA to screen 48’718 plasma samples from 37’894 individuals admitted to almost all clinical departments of the University Hospital Zurich (Fig. 5a-b and Extended data Fig. 8a) for binding to human recPrP_23-230_. We applied stringent criteria: only plasma samples displaying log(EC50) ≥ −2 and logistic-regression fitting error <20% were considered as hits. In a primary high-throughput screen (HTS), 27 individuals (9 females and 18 males) were found to harbor IgG antibodies reacting to human recPrP_23-230_ (Fig. 5c-d). A validation screen confirmed PrP reactivity in the plasma samples of 21 individuals, indicative of a 0.06% prevalence of autoantibody carriers (Fig. 5e). The clinical presentation of individuals with such autoantibodies was heterogeneous (Extended data Table 5), with no statistically significant enrichments found in disease codes (International

Classification of Disease, ICD-10), medication reports, age or sex (Fig.5d, Extended data Fig. 8b,c and Extended data Table 6). None of the patients with anti-PrP reactivity showed signs of prion-related pathology. Repeated longitudinal sampling indicated that 3 individuals sustained their high anti-PrP titres over a time span of several months up to more than a year (Fig. 5g), suggesting that these anti-PrP-autoantibodies are stable over time.

We then assessed the clonality of the anti-PrP-autoantibodies in a subset of PrP-reactive samples. First, we compared by ELISA the λ/κ light-chain ratio in total immunoglobulins vs. anti-PrP autoantibodies. We found that antibodies binding human recPrP_23-230_ have preferential contribution of λ over κ light chains (Fig. 5f). We then purified immunoglobulins from selected plasma samples, confirmed their specific reactivity to human recPrP_23-230_, and assayed them for differential binding to recPrP_23-110_ (FT) versus recPrP_121-230_ (GD). While some patients showed immunoreactivity against the GD, antibodies in other patients targeted both the FT and the GD, suggesting a polyclonal antibody response (Fig. 5h). These findings corroborate the evidence for the existence of naturally occurring antibodies against the prion protein in humans.

## Discussion

Anti-PrP antibodies are effective in cells and mice infected with prions^26, 27^, and may represent a plausible therapeutic strategy^2, 28^. However, although certain anti-PrP antibodies afford neuroprotection to prion-infected mice, others cause extensive neuronal loss^29^ and several antibodies to the GD on PrP result in acute on-target toxicity^11, 21^. Also, antibodies to the OR region of PrP prevent neurotoxicity triggered by GD-binding antibodies and by prions in organotypic slices^11, 22, 30^. Thus, the biological effect of anti-PrP antibodies crucially depends on the targeted PrP epitope. Hence, this study aimed to produce a high-resolution map of neuroprotective epitopes, with the ultimate goal of identifying effective immunotherapeutics. The production of anti-PrP antibodies in wt animals is hindered by self-tolerance^31^. Therefore, anti-PrP monoclonal antibodies have been developed mainly in PrP-deficient mice^17, 32^ or by phage display^33^ and target mostly immunodominant PrP epitopes within the central region and GD of PrP^34^.

Here we screened a synthetic human antibody library by phage display which differs from previous approaches^35, 36^ in several aspects. Firstly, we expanded the number of PrP antigens, instead of using a single PrP fragment for panning, with the intent to discover antibodies to all regions of PrP. Secondly, we deep-sequenced the panning outputs - a strategy that optimizes the detection of extremely rare antibody clones. Finally, we opted to bacterially express antibodies as Fabs which are typically more stable and less susceptible to dimerization than scFv^37^. This enabled us to generate anti-PrP Fabs with highly diverse specificities. In addition, this strategy yielded, besides the 49 Fabs identified by ELISA screening, hundreds of additional rare Fab hits against less antigenic epitopes of PrP.

Having produced a broadly diversified panel of anti-PrP Fabs, we assessed the correlation between their epitope and their biological activity. All OR_51-91_-binding Fabs prevented neurodegeneration in prion-infected COCS. These results are consistent with previous findings^11, 22, 30^ pointing to the OR of PrP as the effector arm of neurotoxicity. In COCS and in prion-infected cells, neuroprotection by OR_51-91_ binders was associated with reduced levels of PK-resistant PrP^Sc^. This effect has not been described for POM2 that blocked the toxic cascade elicited by prions, downstream of its replication. Other OR-targeting antibodies^38^ were described to block PrP^C^ internalization, thus reducing the rate of intracellular conversion to PrP^Sc^. Alternatively, the engagement of the OR by Fab71 and Fab100 may prevent the interaction between PrP^C^ and PrP^Sc^, as suggested for prion-clearing anti-GD antibodies in susceptible cell lines^26, 27^.

Fab83, which binds CC1 (23-KKRPK-27) with high affinity, also afforded neuroprotection. CC1_23-50_-targeting antibodies are rare and, to our knowledge, were never assessed for neuroprotection against prions. However, it is well-established that the CC1 is required for the neurotoxicity of PrP mutants with deletion in the central domain^39^. Transgenic mice expressing PrPΔ23-31 or PrPΔ23-88 not only exhibit prolonged survival after prion inoculation but also accumulate less PrP^Sc^ in their brains^40, 41^. Conversely, Fab83 did not reduce PrP^Sc^ levels in CAD5 cells and in prion infected COCS, yet it was neuroprotective. Furthermore, alanine substitutions of the positively charged amino acids within PrP 23-KKRPK-27 did not prevent PrP^Sc^ formation in prion-infected neuroblastoma cells ScN2a^42^. We conclude that the antibody-mediated neutralization of the CC1 is neuroprotective by other means than arresting PrP^Sc^ generation. We speculate that the CC1 blockade prevents the interaction of PrP with downstream effectors of neurotoxicity, such as the Group-I metabotropic glutamate receptors^43^, which in turn trigger neurotoxicity.

The presence of anti-PrP antibodies in a human repertoire is surprising. Prions do not elicit antibody responses^44^, most likely because of the negative selection of B cells that autoreact to PrP^C^ whose primary amino acid sequence is identical with PrP^Sc 31^. Immunization of wt mice with recombinant PrP did not result in antibodies to N-terminal epitopes, and most antibodies recognized recombinant PrP in ELISA but not native PrP^C^ on cell membranes. We found many Fabs binding within the FT in the synthetic human antibody-phage display library, most of which recognized native PrP on the cell surface. Unexpectedly, sequence comparisons identified PrP-reactive antibodies in published databases of naïve human repertoires from circulating B cells. Finally, we found high-titer PrP autoantibodies in the plasma of unselected hospitalized patients. Certain antibodies to PrP can mimic prion neurotoxicity^11, 22^. At the time of the analysis, none of these subjects had received a diagnosis of prion disease. The clinical presentation of the patients with high-titer PrP autoantibodies was heterogeneous and, although signs of dementia were reported for two of the anti-PrP antibody carriers, we did not find any statistically significant correlation with neurological or any other disease when comparing subjects with and without anti-PrP autoantibodies. Similarly, PrP autoantibody titers present in a subset of *PRNP* mutation carriers neither correlated with the *PRNP* mutation status nor with the onset of clinical prion disease.

The frequency of high-titer anti-PrP antibody carriers (0.06%) is much lower than the occurrence of Fab71-like HCDR3 sequences in published human repertoires. This discrepancy could mean that most anti-PrP specificities exist in a dormant state, or are expressed as B-cell receptors, but do not produce circulating antibodies. It will be interesting to discover the triggers that may ignite antibody production and, possibly, afford protection against prions.

The evidence presented here indicates that the immunological repertoires of unselected humans can contain antibodies against PrP^C^. Such antibodies might protect against prions. The rarity of individuals with high-titer anti-PrP immunity is unsurprising since PrP^C^ is highly expressed in many immune cells including the developing thymus, which in mice results in an almost-insurmountable central tolerance^31^.

We suspect that it is not PrP^C^ that induces adaptive immune responses against prions, but nascent PrP^Sc^ instead. Accordingly, clinically silent prion generation may occasionally occur in healthy individuals^46^. PrP^Sc^ aggregates arising de novo may result in exposure of neoepitopes and/or epitopes occluded in cell-borne PrP^C^. The resulting immune response may clear any nascent prions, akin to the immune surveillance against neoplastic cells. Progressive senescence of adaptive immunity^47^, which is well-documented for a variety of infectious diseases, may explain why both sporadic and familial prion diseases flare up mostly in late life.

In conclusion, the generation of antibodies to the whole PrP epitope space provides new tools to understand the mechanism of neurodegeneration conveyed by prions. The presence of prion protein binders in human antibody repertoires and of anti-PrP reactivity in human plasma points to a potential source of immunotherapeutics against prion diseases.

## Methods

### Experimentation with human samples

All experiments and analyses involving samples from human donors were conducted with the approval of the local ethics committee (KEK-ZH-Nr. 2015-0561 and BASEC-Nr. 2018-01042) and in accordance with the provisions of the Declaration of Helsinki and the Good Clinical Practice guidelines of the International Conference on Harmonisation.

### Animal experiments

All animal experiments were conducted in strict accordance with the Swiss Animal Protection law (*Tierschutzgesetz* and *Tierschutzverordnung*) of the *Swiss Bundesamt für Lebensmittelsicherheit und Veterinärwesen BLV*. The Animal Welfare Committee of the Canton of Zurich approved all animal protocols and experiments performed in this study (animal permits Versuchstierhaltung 123, ZH90/2013, ZH139/16).

### Antigen production

Synthetic PrP peptides CC1_23-50_, N-OR_39-66_, F-OR_51-91_ and CC2-HC_92-120_ were synthesized (EZbiolabs) with a convenient C-terminal biotin moiety, separated by a short linker. Recombinant mouse and human PrP proteins were produced as described^48^.

### Construction of the synthetic human Fab phage library

A synthetic human Fab phagemid library (Novartis Institutes for BioMedical Research) was used for the phage display selections. A gene fragment encoding the germlines frameworks combinations IGHV3-23 and IGKV1-39, IGHV3-23 and IGLV3-9, IGHV3-30 and IGKV3-15, IGHV3-15 and IGLV1-47, were synthesized by Invitrogen’s GeneArt service in Fab format and cloned into a phagemid vector serving as the base templates. These human germlines were used as they display favorable frameworks combinations for a phage display library^49^. The phagemid vector consists of Ampicillin resistance, ColE1 origin, M13 origin and a bi-cistronic expression cassette under a lac promotor with OmpA - light chain followed by PhoA–heavy chain – Flag – 6xHis – Amber stop – truncated pIII (amino acids 231 – 406).

Only HCDR3 was diversified and primers were designed to incorporate up to 11 amino acids at defined ratios mimicking their natural occurrences: aspartic acid, glutamic acid, arginine, histidine, serine, glycine, alanine, proline, valine, tyrosine, tryptophane. Leucine and phenylalanine were also allowed at a certain position of HCDR3. Certain residues were omitted on purpose to remove potential post translational modification sites. Randomized primer synthesis was performed using the Trinucleotide technology (ELLA Biotech) in order to exclude stop codons, methionine, cysteine and asparagine.

Lengths between 8, 10, 12, 14, 16 and 20 amino acids were allowed, in which the last two amino acids were kept constant with the sequence Asp-Tyr for length 8 to 16 and Asp-Val for length 20. The design of the final two HCDR3 amino acids reflects human VDJ recombination. Short HCDR3s more often use J-fragment IGHJ4 with “DY” at the end of HCDR3, while longer HCDR3s (here 20 aa) more often use IGHJ6 with “DV” at the end of HCDR3.

Library inserts were generated by PCR using Phusion High Fidelity DNA polymerase (NEB Biolabs). The resulting HCDR3 library inserts were ligated into the base templates, transformed into E.coli TG1F+ DUO (Lucigen) with a minimal library size of 3E+08 transformants per HCDR3 length and phages were produced using VCSM13 helper phage (Agilent Technologies) using standard protocols.

### Phage display for isolation of PrP binders

Depending on the HCDR3 length, the library was divided into two sub-pools for panning: short (8-10 aa) and long (12-20 aa) HCDR3 - that were run in parallel for the selection. First, two rounds of selection were performed by coating 96-well Maxisorp plates (Nunc) with decreasing amount of mouse recPrP_23-231_ (1 µM and 0.5 µM respectively, in PBS), overnight at 4°C. PrP-coated plates were washed and blocked with Superblock for 2 h. Input of 4 x 10^11^ phages in 300 µl of PBS was used for the first round of panning. After 2 h blocking with Chemiblocker (Millipore), the phages were incubated with recPrP_23-231_-coated wells for 2 h at room temperature (RT). The non-binding phages were then removed by extensive washing with PBS 0.05% Tween-20 (PBS-T) while mouse recPrP_23-231_ bound phages were eluted with of 0.1 M Glycine/HCl, pH 2.0 for 10 min at RT and the pH was neutralized by 1M Tris pH 8.0. Eluted phages were used to infect exponentially growing Amber suppressor TG1F+ cells (Lubio Science). Infected bacteria were cultured in 2YT/Carbenicillin/1% glucose medium overnight at 37°C, 200 rpm and superinfected with VCSM13 helper phages. The production of phage particles was then induced by culturing the superinfected bacteria in 2YT/Carbenicillin/Kanamycin medium containing 0.25 mM isopropyl β-D-1-thiogalactopyranoside (IPTG), overnight at 22°C, 180 rpm. Supernatant containing phages from the overnight culture was used for the second panning round. Output phages from the second round were purified by PEG/NaCl precipitation, titrated, and selected in the following third rounds. For the third rounds either mouse recPrP_23-231_, recPrP fragments were used as bait at 0.25 uM in PBS for coating 96-well Maxisorp plates. Selection for binders to biotinylated PrP peptide was performed with the antigen in solution followed by capture on neutravidin-coated wells. Selection for CC1_23-50_ binders was also performed in solid phase by capturing of CC1_23-50_ biotinylated peptide on neutravidin coated wells. After the third round of selection, DNA minipreps were prepared from the panning output pools by QIAprep Spin Miniprep kit (Qiagen) and the whole anti-PrP Fab enriched library was either used for NGS analysis or used for bacteria transformation.

### ELISA screening of bacterially expressed Fab-phagemid vectors

Electrocompetent non-Amber suppressor MC1061 bacteria (Lubio Science) were transformed with pPD2-Fab phagemid vectors to perform the primary ELISA screening. DNA used for transformation derived from the following panning sets: full-length recPrP_23-231_, CC1_23-50_ in solid phase and in liquid phase, N-OR_39-66_, F-OR_51-91_ and CC2-HC_92-120_. Single colonies were picked randomly by an automated colony picker and cultured in 384 well plate (Nunc) in 2YT/Ampicillin/1% glucose medium over night at 37°C, 80% humidity, 500 rpm. These precultures were used to prepare glycerol stock master plates. Expression plates were prepared by inoculating 2YT/Carbenicillin/0.1% glucose medium, followed by induction with 1mM IPTG. After overnight culture at 25°C, 80% humidity, bacteria cultures were lysed for 1.5 19 h at 400 rpm, 22°C in borate buffered saline pH 8.2 containing EDTA-free protease inhibitor cocktail, 2.5 mg/ml lysozyme and 40 U/ml benzonase. Fab-containing bacteria lysate was blocked with Superblock for 2 h and used for the ELISA screening. The following antigens were coated on separate 384-well ELISA plates: anti-Fd antibody (The Binding Site GmbH) 1:1000 in PBS, to check the expression level of each Fab clone in bacteria lysates, and mouse recPrP_23-231_ at 87 nM in PBS. Antigen-coated ELISA plates were washed twice with TBS-T and blocked with Superblock for 2 h. Fab containing bacteria lysates from the expression plate were transferred to corresponding wells of the ELISA plates. Wells containing only medium or POM antibodies at 50 nM in TBS-T were included as background and positive controls, respectively. After 2 h, ELISA plates were washed three times with TBS-T and anti-human F(ab’)_2_-alkaline phosphatase conjugated antibody (1:5000 in TBS-T) was added. After 1h incubation at RT, followed by three washings with TBS-T, Attophos substrate (Roche) was added and, after 10 min incubation, the ELISA signal (fluorescence) was measured. Fabs from bacteria lysates producing an ELISA signal 5-10 times higher than the technical background, which was calculated as the average of the coated well containing only medium, were considered as mouse recPrP_23-231_ binder candidates. All the identified hits were checked in confirmatory ELISA screening. Anti-CC1_23-50_ anti-OR_51-91_, anti CC2-HC_92-120_ and anti-GD binder hits were identified. Corresponding Fabs were subcloned into the Fab expression vector pPE2. pPE2-Fab plasmids were transformed in TG1F-chemicompetent cells and grown on LBagar/Kanamycin/1% glucose plates. Bacteria cultures of the selected clones were used for DNA minipreps followed by Sanger sequencing using the following sequencing primers: VH (5’-GATAAGCATGCGTAGGAGAAA-3’) and M13Rev (5’ - CAG GAA ACA GCT ATG AC – 3’).

### Affinity maturation of selected Fab3 and Fab71

Fab3 and Fab71 were further engineered to improve their affinity by using affinity maturation cassette libraries (Novartis) with either diversification in the HCDR2 or in the LCDR3. The HCDR2 and the LCDR3 sequences repertoires were diversified according to naturally occurring repertoire of rearranged human CDR sequences.

For LCDR3 libraries, primers were designed to incorporate up to 11 amino acids at defined ratios mimicking their natural occurrences: aspartic acid, glutamic acid, arginine, histidine, threonine, serine, glycine, alanine, leucine, valine, tyrosine. Glutamine, proline and tryptophan were also allowed at certain positions of LCDR3. LCDR3 lengths of 9 and 10 amino acids were allowed for IGKV1-39 and IGKV3-15, in which the last threonine was kept constant.

For HCDR2 libraries, primers were designed to incorporate up to 10 amino acids at defined ratios mimicking their natural occurrences: aspartic acid, glutamic acid, arginine, histidine, threonine, serine, glycine, alanine, valine, tyrosine. Isoleucine, and tryptophan were also allowed at certain positions of HCDR3. A length of 17 amino acids was allowed for the HCDR2 of IGHV3-23 and IGHV3-30, in which the last seven amino acids were kept constant (YADSVKG).

For LCDR3 and HCDR2 maturation cassettes, the synthesis of randomized primer was performed using the Trinucleotide technology (ELLA Biotech) in order to exclude stop codons, methionine, cysteine and asparagine. The randomization of LCDR3 and HCDR2 was performed by two PCR steps using Phusion High Fidelity DNA polymerase (NEB Biolabs). A first randomization PCR was performed using randomized primers (a forward primer to randomize LCDR3 and a reverse primer to randomize HCDR2). In a second step the randomized fragments are amplified by using primers which also introduce restriction sites at both ends. This amplification PCR was then ligated into the base templates phagemid vectors using the restriction sites, transformed into E.coli TG1F+ DUO (Lucigen) with a minimal library size of 1E+08 transformants. For the HCDR2 affinity maturation libraries of Fab3 and Fab71, MfeI-HF and BssHII restriction digestion was performed to replace the parental HCDR2 sequence with randomized cassettes of diversified HCDR2 sequences. For the LCDR3 affinity maturation libraries of Fab3 and Fab71, MfeI and EcoRI restriction digested parental HCDR3 was grafted into the affinity maturation phagemid library with randomized LCDR3. HCDR2 and LCDR3 phagemid libraries of Fab71 and Fab3 were transformed into Top10F’ competent cells (Invitrogen), phage particles were produced by superinfection with VCSM13 helper phage and then precipitated by PEG/NaCl. These four phage libraries (Fab3 and Fab71 HCDR2; Fab3 and Fab71 LCDR3) were subjected to three rounds of panning by alternating exposure to mouse recPrP_23-231_ and to the respective biotinylated peptides containing the targeted epitope (CC1_23-50_ -biotinylated peptide for Fab3 and OR-biotinylated for Fab71). To drive the selection for high affinity binders, two strategies were used. As a standard condition for maturation, washing stringency was increased as compared to the panning conditions used for selection of the parental Fabs and the amount of antigen from one round to the next was decreased (from 200 nM at the first round to 1 nM at the third round). As an alternative approach, panning was performed in competition with 5-fold molar excess of the parental Fabs. After the third round of panning, DNA minipreps were used for NGS screening. Moreover, reading frames of the enriched pools of matured Fab3 and Fab71 were inserted into pPE2 vector for expression in bacteria, followed by ELISA screening. For each Fab, 4 output pools were obtained. For each pool, 368 colonies were randomly picked and transferred to a 384-well plate for primary ELISA screening using bacteria lysates containing soluble Fabs, as described above. To differentiate matured Fabs with higher affinity as compared to the parental version, off-rate and on-rate ELISA screens were performed using mouse recPrP_23-231_ coated ELISA 384-well plates. Briefly, for off-rate selection, Fab containing E. coli lysates were exposed for 2 h to recPrP_23-231_ coated wells and, after ten washing steps lasting from 1 h to over-night, anti-human F(ab’)_2_-alkaline phosphatase conjugated antibody was added, followed by Attophos substrate (Roche) and measurement of the ELISA signal (fluorescence) as described above. During the on-rate ELISA screen, the incubation time of the diluted Fab containing E. coli lysates with recPrP_23-231_ coated wells was only 20 minutes, followed by 3 fast washes. From the primary screening 95 positive clones were sequenced and assessed in confirmatory screens by off-rate and on-rate ELISA whereby retention of epitope specificity was checked by measuring the binding of affinity matured versions of Fab3 and Fab71 to CC1_23-50_ and F-OR_51-90_ biotinylated peptides. Hits consisting of 19 Fab3 matured clones and 5 Fab71 matured clones were selected and purified by IMAC.

### Sample preparation for NGS

Polyclonal DNA minipreps isolated from the third panning output pools were used as PCR template to amplify the HCDR3 of the selected Fabs and add the adapters required for sequencing on Illumina sequencer MiSeq. The PCR protocol has been described^50^.

The PCR product was purified and DNA concentration was measured using the Qubit DNA High sensitivity kit (Invitrogen). Samples were analyzed on a MiSeq using MiSeq reagent kit *MiSeq v2 Reagent kit 300 cycles PE*.

### NGS data analysis

The data analysis of the NGS FastQ output files was performed as described^50^. For each panning output, 100’000 sequences were analyzed using the fixed flanking sequences on the boundary of variable region as template to locate and segment out the HCDR3 sequence. ∼40’000 to 70’000 HCDR3 sequences were identified depending on the panning output pools, and included into frequency reports in CSV format.

For determination of clones to high immunogenic PrP epitopes, we selected HCDR3 displaying ≥ 20 NGS counts in recPrP_23-231_ panning in 100’000 analyzed sequences. For rare clones against less immunogenic PrP epitopes, HCDR3 were identified according to the following criteria: NGS count in recPrP_23-231_ panning = 1 and, to avoid selecting for sequences resulting from PCR or sequencing errors, sum of the NGS counts across all the panning outputs ≥ 10.

### Rescue of clones identified in NGS

Fab clones of interest based on the NGS binding profile can be retrieved from the polyclonal DNA output after phage panning by an assembly PCR approach. For FabRTV, first two separate PCR reactions were executed to amplify the Fab in two fragments, one ranging from VL until HCDR3 (LC –HCDR3 PCR) and the second one ranging from HCDR3 until the tag (HCDR3-CH1-tag). Primers specific for FabRTV HCDR3 sequence were used (FabRTV_HCDR3_fw: 5’-TCGTTACGTTCGTGGTTACGGTTCTCC-3’; FabRTV_HCDR3_rw: 5’-GGAGAACCGTAACCACGAACGTAACGA-3’ in combination with PAL_fw (PAL_fw: 5’-GGAAACAGCTATGACCATGATTACGCCAAG-3’ and Flag_rw (Flag_rw: 5’-CGCACCTTTGTCATCGTCATCTTTATAGTCG-3’). The polyclonal DNA pool from the third round was taken as template. LC–HCDR3 and HCDR3-CH1-tag amplicons were generated using Phusion High-Fidelity DNA Polymerase and the following conditions: 98°C for 30 s, 98°C for 10 s, 62-65°C for 30 s, 30 cycles at 72°C for 100 s (LC –HCDR3 PCR) or 72°C for 40 s (HCDR3-CH1-tag), and 72°C for 5 min. PCR products were purified. LC –HCDR3 and HCDR3-CH1-Flag fragments were then assembled in the next PCR using the primers PAL_fw and Flag_rw. The assembly PCR was performed using 60 ng LC-HCDR3 and 20 ng HCDR3-CH1-Flag as template in two steps. A first PCR sequence was run without primers (98°C for 30 s, 98°C for 10 s, 55°C for 30 s, 5 cycles at 72°C for 100 s, and 72°C for 7 min). Then, primers PAL-fw and Flag-rw were added and the second PCR step was run (98°C for 30 s, 98°C for 10 s, 60°C for 30 s, 25 cycles at 72°C for 120 s, and 72°C for 7 min). The assembly PCR product was purified, digested by NruI and EcoRI and ligated into pPE2 expression plasmid. Retrieved clones were transformed into XL1-Blue electrocompetent cells and DNA was prepared from isolated colonies for Sanger sequencing.

### Expression and purification of selected anti-PrP Fabs

Chemical competent BL21(DE3) cells (Invitrogen) were transformed with selected pPE2-Fab plasmids and grown on LBagar/Kanamycin/1% glucose plates. A single colony was inoculated into 20 ml of 2xYT/Kanamycin/1% glucose pre-culture medium and incubated for at least 4 h at 37°C, 220 rpm. One liter of 2YT-medium containing Kanamycin/0.1% glucose was inoculated with 20 ml pre-culture and Fab expression was induced by 0.75 mM IPTG followed by incubation over night at 25°C, 180 rpm. The overnight culture was centrifuged at 4000 x g at 4°C for 30 min and the pellet was frozen at −20°C. For Fab purification, thawed pellet was resuspended into 20 ml lysis buffer: 0.025 M Tris pH 8; 0.5 M NaCl; 2 mM MgCl_2_; 100 U/ml Benzonase (Merck); 0.25 mg/ml lysozyme (Roche), EDTA-free protease inhibitor (Roche), and incubated for 1 h at RT at 50 rpm. Lysate was centrifuged at 16000 x g at 4°C for 30 min and supernatant was filtrated through 0.22 μM Millipore Express®Plus Membrane. Fab purification was achieved via the His6-Tag of the heavy chain by IMAC. Briefly, after equilibration of Ni-NTA column with running buffer (20 mM Na-phosphate buffer, 500 mM NaCl, 10 mM Imidazole, pH 7.4), the bacteria lysate was loaded and washed with washing buffer (20 mM Na-phosphate buffer, 500 mM NaCl, 20 mM Imidazole, pH7.4). The Fab was eluted with elution buffer (20 mM Na-phosphate buffer, 500 mM NaCl, 250 mM Imidazole, pH7.4). Buffer exchange in PBS was performed using PD-10 columns, Sephadex G-25M (Sigma).

### ELISA for epitope specificity of purified Fabs

For epitope profiling of the Fabs by ELISA, 384-well SpectraPlates (Perkin Elmer) were coated with the following antigens at 87 nM in PBS, at 4°C overnight: mouse recPrP_23-231_, recPrP_23-110_, recPrP_90-231_, recPrP_121-231_, BSA and neutravidin. Plates were washed three times in PBS-T and blocked with 50 µl per well of Superblock for 2 h at RT. Biotinylated PrP peptides CC1_23-50_, N- OR_39-66_; F-OR_51-91_ and CC2-HC_92-120_ were captured on neutravidin-coated wells for 1h at RT. Purified Fabs at 150nM in PBS-T were then incubated for 2 h at RT. After three washing with PBS-T, binding Fabs were detected by anti-human F(ab’)_2_-alkaline phosphatase conjugated antibody (1:1000 in PBS-T). After 1h incubation at RT, followed by three washings with PBS-T, Attophos substrate (Roche) was added and, after 10 min incubation, the ELISA signal was measured. The relative affinities of Fab binding to PrP were determined by using the Fabs at serial dilutions by measuring the concentration of Fab required to achieve 50% maximal binding (EC50). Data analysis was performed using Non-linear regression (GraphPad Prism, GraphPad Software).

### Competition ELISA for epitope mapping

The approach described by Polymenidou et at.^17^ was used. A library of overlapping 12-mer mouse PrP-peptides, shifted by two amino acids, was synthesized by Jerini, Germany. It consisted of 50 peptides, spanning the whole FT of the mouse PrP sequence. Mouse recPrP_23-231_ at 65 nM in PBS was coated to 384 well HB plates (Perkin Elmer), overnight at 4°C. Plates were washed three times with PBS-T and blocked with Superblock for 2 h at RT.

After washing, plates were incubated with 400 nM of anti-PrP Fabs in 1% Superblock in PBST, with or without 175 molar excess of each dodecameric peptide (final concentration of 70 uM). After 2 h at RT plates were washed three times and incubated with anti-human Fab specific-HRP conjugated secondary antibody (Sigma) for 1 h at RT. TMB, was added, incubated for three minutes at RT, and the chromogenic reaction stopped by addition of 0.5 M H_2_SO_4_. The absorbance at 450 nm was measured in a plate reader (Perkin Elmer, EnVision).

### ELISA with dilution series for EC50 determination

To compare affinity matured and parental Fabs, ELISA 384 well HB plates (Perkin Elmer), were coated with mouse recPrP_23-231_ at 87 nM in PBS overnight at 4°C. After three washing steps in PBS-T and blocking with Superblock for 2 h at RT, plates were incubated for 2 h at RT with 1:2 serially diluted Fabs at starting concentration of 150 nM in PBS-T. Plates were incubated with anti-human Fab specific-HRP conjugated secondary antibody (Sigma) for 1 h at RT, followed by washing and development of chromogenic reaction by TMB, as described above. The absorbance at 450 nm was measured in a plate reader (Perkin Elmer, EnVision). For Fabs that were identified by mining of NGS datasets of human antibody repertoires, human recPrP_23-230_ at 87 nM in PBS was used for coating and serially diluted Fabs were tested starting from 3 µM in PBS-T.

### Measurement of binding kinetics by surface plasmon resonance (SPR)

Binding kinetic of the purified Fabs to full length PrP was monitored at 25 °C using ProteOn XPR36 surface plasmon resonance biosensor (Bio-Rad Laboratories). Mouse recPrP_23–231_ was immobilized on a carboxymethylated-dextran sensor chip (CM5, Bio-Rad) by EDC/NHS chemistry at either low or high density (1000 and 100 resonance units (RU), respectively) in the test flow cells. Two flow cells were left untreated, one was exposed to activation and deactivation buffers used in EDC/NHS chemistry and one was used to immobilize BSA as a control. Six serial 1:2 dilution of Fab at starting concentration of 100 nM were prepared in SPR running buffer (10 mM Tris/HCl buffer pH 8.5, 500 mM NaCl, 3 mM EDTA, 0.05% Tween-20) and injected at flow rate of 100 µl/min for 120 s. Association and dissociation were monitored over a 300 s interval. BSA immobilized flow cell was used as negative control for the antigen and responses were corrected by referring to the flow cell where only SPR running buffer was injected. The sensor chip was regenerated after each round of Fab injection by flowing the sensor chip chambers with 20 nM NaOH for 60 s at 25 ul/min. Sensorgrams were fitted by Langumir equation (the 1:1 interaction model) and antibody binding constants (*kon* and *koff* and *KD*) were calculated by the Bio-Rad ProteOn manager software.

### Epitope confirmation

SPR approach was used to confirm the epitope specificity of the different anti-PrP Fabs. An NLC sensor chip (Bio-rad) was used to immobilize the biotinylated PrP peptides CC1_23-50_, N- OR_39-66_, FL-OR_51-91_ and CC2-HC_92-120_ in different flow cells at 700 RU. Dilution series of each Fab were injected at 100 µl/min starting from 100 nM and real time binding was monitored (120 s association time and 300 s dissociation time, regeneration by 20 nM NaOH for 60 s at 25 µl/min).

### Establishment of CAD5 Prnp^-/-^ cells by CRISPR

The CAD5 cell line derived from Cath.a-differentiated (CAD) cells was reported to be responsive to several prion strains^51^. For control in our experiments, CAD5 *Prnp^-/-^* cells were established by CRISPR/Cas9-mediated gene ablation. Briefly, a guide RNA (gRNA) targeting exon 3 of the *Prnp* gene (5′-TCA GTC ATC ATG GCG AAC CT −3′) was cloned into the pKLV-U6gRNA(BbsI)-PGKpuro2ABFP vector (Addgene 50946,) to generate pKLV-*Prnp* sgRNA plasmid. A CAD5 cell line stably expressing Cas9 (CAD5-Cas9) was generated by transfecting linearized pCMV-hCas9 (Addgene 41815,). CAD5-Cas9 cells were then transfected with pKLV-*Prnp* sgRNA by Lipofectamine 2000 (Invitrogen, 11668-019). Transfected cells were selected and enriched by puromycin (2 μg/ml) and single cell clones were obtained by limited dilution. Cell clones were grown and expanded. To confirm gene editing of the *Prnp* gene in cell clones, a region embracing gRNA was PCR-amplified using a forward primer (5′-TGC AGG TGA CTT TCT GCA TTC TGG-3′) and reverse primer (5′-GCT GGG CTT GTT CCA CTG ATT ATG GGT AC-3′), the PCR product was cloned into pCR-Blunt II-TOPO vector (Invitrogen, 45-0245) and sequenced to verify the frameshifted indels. To confirm gene editing of the *Prnp* gene in cell clones, a region embracing gRNA was PCR-amplified using a forward primer (5′-TGC AGG TGA CTT TCT GCA TTC TGG-3′) and reverse primer (5′-GCT GGG CTT GTT CCA CTG ATT ATG GGT AC-3′), the PCR product was cloned into pCR-Blunt II-TOPO vector (Invitrogen, 45-0245) and sequenced to verify the frameshifted indels.

### Flow cytometry

Binding of Fabs to wtPrP^C^ on the surface of CAD5 PrP^+/+^ cells was determined by flow cytometry. CRISPR/Cas9-derived CAD5 *Prnp^-/-^* cells were used as negative control. Briefly, Confluent CAD5 *Prnp^+/+^* and CAD5 *Prnp^-/-^* cells in 75-cm flasks were washed with PBS, detached in dissociation buffer (2 mM EDTA, PBS) and washed twice in ice-cold FACS buffer (10 mM EDTA, 2% FBS in PBS) by centrifugation at 190 x g for 5 minutes at 4 °C. Cells were stained with Trypan Blue, counted and density of live cells adjusted to 0.5 × 10^6^ cell in 100 µl. 100 µl of cell suspension were added to each well of a 96-well round bottom tissue culture plate, and centrifuged at 800 x g for 2 minutes at 4 °C. Cells were incubated for 30 minutes at 4 °C with the Fabs at 150 µg/ml. After washing twice in FACS buffer, Fabs bound to cell surface wtPrP^C^ was detected by incubating the cells with APC-labeled anti-human Fab secondary antibody (Jackson) diluted 1:400 for 30 minutes at 4 °C in the dark. Cells stained with anti-PrP POM1 Fab and APC-labeled anti-mouse Fab were used as a positive control while staining with secondary antibodies was used to establish the background fluorescence. Cells were washed 3 times with ice-cold FACS buffer and transferred into micro-FACS tubes. Data were acquired using FACSCanto II cytometer (BD Biosciences, San Jose, CA, USA) and the geometric mean fluorescence (MFI) of stained cells was measured using FlowJo software.

### Immunohistochemistry

Staining was performed on sections from brain tissues fixed in formalin and embedded in paraffin. After deparaffinization through graded alcohols and heat-induced antigen retrieval in citrate buffer (0.01 M; pH 6), sections were blocked and incubated with the anti-PrP Fabs at 12 µg/ml and antibody for Microtubule Associated Protein 2 (MAP2) (1∶500, Abcam). Double staining was performed with fluorescently-labeled secondary antibodies (goat anti Human IgG F(ab’)2:Tritc 1:150, Bio-Rad and goat anti-rabbit 1:500, Alexa Fluor 488, Invitrogen), followed by nuclear staining by DAPI (Life technologies). Images acquisition was done by using the fluorescence microscope (BX-61; Olympus), equipped with a cooled black/white charge-coupled device camera, using identical acquisition settings. Images were analyzed using the image-processing software CellF.

### Immunoprecipitation

For immunoprecipitation of PrP from brain extracts, the tissue was homogenized in ice-cold IP buffer (75 mM NaCl, 1% Igepal, protease inhibitor mixture (Sigma), 50 mM Tris-Cl, pH 7.4). After centrifugation at 1000 x g for 5 min, the supernatant was recovered and protein content quantified by BCA assay (Pierce). 1 ml of Dynal sheep-anti mouse IgG paramagnetic beads were non-covalently coupled with 60 µg of anti-His mAb (Invitrogen) in PBS plus 0.1% immunoglobulin-free BSA (Sigma) for 1 h at RT on a rotating wheel. Three molar excess of the His-tagged Fab were added to allow the formation of the complex with the anti-His antibody-coupled dynabeads and after 1 h incubation, three washes in PBS plus 0.1% immunoglobulin-free BSA were performed. 500 µg of brain homogenate was diluted in IP buffer and incubated first with 20 µl of dynabeads coupled only to the anti-His antibody without the Fab to remove the unspecific binding of PrP to the beads and to the IgG. Then the cleared brain homogenate was incubated with 50 µl of Fab-anti-His antibody-coupled dynabeads and immunoprecipitation was performed for 2 h at RT on a rotating wheel. After five washes with 150 mM NaCl, 0.5% Igepal, 50 mM Tris-Cl, pH 7.4, elution of immunoprecipitated PrP was performed by incubation for 3 h at 4°C with 62 µg of 12-mer PrP peptides in 40 µl of PBS plus protease inhibitors. Peptide P1: KKRPKPGGWNTG was used for competition with Fab3 and peptide P17 TWGQPHGGGWGQ was used for competition with Fab71 and Fab100. Peptides P2 RPKPGGWNTGGS and P15 H-PQGGTWGQPHGG-OH were used as negative controls. Eluate was finally collected and supplemented with loading buffer (NuPAGE, Invitrogen) for western blot analysis using the mouse monoclonal antibody POM1 and the rabbit polyclonal antibody XN (produced in house) for PrP detection.

### Immunoblot analysis

For epitope confirmation of the Fabs by western blot, brain from transgenic mice expressing different PrP deletion mutants was homogenized in 10 volumes of lysis buffer (50 mM Tris-HCl pH 8, 0.5% Na deoxycholate, and 0.5% Igepal, protease inhibitors (complete Mini, Roche) using TissueLyser LT for 5 min for 2 cycles. After centrifugation at 1000 x g for 5 min at 4°C to remove debris, protein concentration in the post nuclear supernatant was measured by BCA. 25 µg of total proteins were separated by SDS-PAGE (Novex NuPAGE 12% Bis-Tris Gels) and transferred to PVDF membrane. Membranes were blocked with 5% milk in TBS-T for 1h at RT and incubated overnight at 4°C with anti-PrP Fabs at 10 µg/ml. After washing, the blots were incubated with secondary antibody horseradish peroxidase (HRP)-conjugated goat anti-human Fab IgG (H+L) (Sigma, A0293) diluted 1:10000 in blocking buffer for 1 h at RT. Blots were developed using Luminata Crescendo Western HRP substrate (Millipore) and visualized using the Stella system (model 3200, Raytest).

For PrP^sc^ detection in prion infected cultured organotypic cerebellar slices (COCS), cerebellar slices were washed in PBS and then scraped off the membrane using 1 ml of ice cold PBS, pelleted and homogenized by trituration in lysis buffer. Samples were centrifuged at 1000 x g for 5 min at 4°C and protein concentration in the post nuclear supernatant determined by BCA. Samples were adjusted to 20 µg protein in 20 µl and digested with 5 µg/ml proteinase-K (PK) in digestion buffer (0.5% wt/vol sodium deoxycholate and 0.5% vol/vol Nonidet P-40 in PBS) for 30 min at 37°C. PK digestion was stopped by adding loading buffer (NuPAGE, Invitrogen) and boiling samples at 95°C for 5 min. Proteins were separated on a 12% Bis-Tris polyacrylamide gel and blotted onto a PVDF membrane by using the iblot apparatus (Bio-rad). Membranes were blocked with 5% top-block in PBS-T followed by incubation with POM1 mouse IgG_1_ antibody (200 ng ml^−1^). After washing, secondary antibody rabbit anti–mouse IgG_1_ (1∶10,000, Zymed) used was. Blots were developed Luminata Crescendo Western HRP substrate (Millipore) and visualized using the Fuji Stella (Bio-Rad).

#### Antibody treatment in cultured organotypic cerebellar slices

Cultured organotypic cerebellar slices were prepared from 9-12 day old *tg*a20 pups as previously described ^52^. For prion experiments, COCS were infected as free-floating sections with 100 µg per 10 slices of RML6 (Rocky Mountain Laboratory strain mouse-adapted scrapie prions at 6 passage) brain homogenate from terminally sick prion-infected mice. As control, non-infectious brain homogenate (NBH) from CD1-inoculated mice was used. After incubation with brain homogenates diluted in physiological Grey’s balanced salt solution for 1 h at 4°C, the slices were washed and 5-8 sections were seeded on a 6-well PTFE membrane insert. Treatment with Fab (550 nM) was started 1 day after plating and supplied at every medium exchange. At 45 days in culture, slices were fixed and processed for immunocytochemistry.

#### Immunofluorescence staining of COCS and NeuN morphometry

For immunofluorescence staining, COCS were washed twice in PBS and fixed in 4% formalin overnight at 4°C. After washing in PBS, the COCS were incubated with blocking buffer (0.05% Triton X-100 vol/vol, 0.3% goat serum vol/vol in PBS) for 1 h at RT and then incubated for 3 days at 4°C with the monoclonal mouse anti-NeuN antibody conjugated with Alexa-488 (clone A60, Life Technologies) at a concentration of 0.5 µg mL^-1^ into blocking buffer. Slices were then washed two times with PBS for 15 min and then incubated with 4,6-diamidino-2-phenylindole (DAPI) (1 µg mL^-1^) in PBS at RT for 30 min to visualize cell nuclei. Two subsequent washes in PBS were performed and COCS were mounted with fluorescence mounting medium (DAKO) on glass slides. NeuN morphometry was performed by image acquisition on a fluorescence microscope (BX-61, Olympus) at identical exposure times. Morphometric analysis was performed to quantify the area of immunoreactivity on unprocessed images using a custom written script for cell^P (Olympus).

#### Prion-infected CAD5 cells

CAD5 *Prnp^+/+^* and *CAD5 Prnp^-/-^* cells were cultured in phenol red free OPTI-MEM supplemented with 10% FBS, Glutamax, penicillin G and streptomycin at 37°C in 5% CO_2_/95% air. CAD5 *Prnp^+/+^* and CAD5 *Prnp^-/-^* cells (5 × 10^4^ in 2 ml of medium) were seeded into 6-well plates (Corning Costar) and cultured for 1–2 days before exposure to 500 µg/ml of prion-infected mouse brain homogenate. Non-infectious brain homogenate (NBH) from CD1 mice at the same dilution was used as control. Treatment with Fabs at 10 µg/ml (200 nM) was initiated 2h after infection and repeated at every split by spiking into the culture medium. Three biological replicates were prepared for each condition. The inoculum was removed after 3 days and the cells were split 1:5 every 3–4 days. After 4 splits (14 days in vitro, DIV) the cells were assayed for PrP^Sc^ by the TR-FRET assay as described below.

#### Quantification of PrP^Sc^ by homogeneous-phase TR-FRET

Prion infected CAD5 *Prnp^+/+^* and CAD5 *Prnp^-/-^* cells, and NBH-inoculated control cells, at 14 DIV in 6-well plates, were detached by pipetting and 10^4^ cells in 40 µl of medium were seeded per well of a 384-well plated (Perkin Elmer) and cultured for one additional day. To detect PrP^Sc^ selectively in homogeneous phase, all the following steps were performed by sequentially adding the reagents into each well without any washing step. First, proteinase-K (PK) digestion was carried out by adding 10 µl of lysis buffer containing 50 or 25 µg/ml of PK (10 or 5 µg/ml PK final concentration) and plates were incubated at 37°C for 1.5 h at 1000 rpm. PK digestion was stopped by adding 4 µl/well of PMSF (final concentration 2 mM) and incubated for 10 min at RT. A denaturation step was performed by NaOH at 58 mM for 10 min at RT to enhance the accessibility of buried epitopes in PrP^Sc^. After neutralization by NaH_2_PO_4_ buffer (58 mM, pH 4.3) for 10 min at RT, plates were processed for FRET measurement. To detect PK-resistant PrP^Sc^, anti-PrP holo-antibodies POM19 and POM1 recognizing two close epitopes in the globular domain^17^ were used. As described^53^, first, 5 µl/well (2.5 nM final) POM19 (made in-house), coupled to Europium (Eu), and diluted in 1X Lance Detection Buffer (Perkin Elmer) was dispensed. Then, 5 µl (5 nM final) POM1 conjugated to Allophycocyanin (APC) was added. TR-FRET readout was performed after 1 h incubation at 4°C using an EnVision 2105 Multimode Plate Reader (PerkinElmer) with previously defined measurement parameters^53^. The collected fluorescence data were corrected by both background and spectral overlap between Eu and APC channel. Net FRET calculations and blank subtractions were performed as previously described^53^.

#### Identification and cloning of Fabs from human antibody repertoire datasets

NGS datasets of antibody repertoires from published human healthy donors, here referred to as DW, BB and DK ^23–25^. Sequencing data included naïve and memory B-cells of 14 donors in total that could be downloaded already pre-processed. HCDR3 amino acid sequences harboring the SYGGY sequence of Fab71 were analyzed and selected if the overall difference to Fab71 HCDR3 was ≤ 3 residues. Afterwards, HCDR3 duplicates, with the same VH, DH and JH segments, found in technical replicates were removed. Then, unique Fab71 similar HCDR3 in each donor were listed and compared across all the subjects from all the datasets. This identified HCDR3 similar to Fab71 that were present in more than a donor with different VH, DH and JH segments. Nucleotide sequences of the variable regions, including each selected HCDR3 Fab71 similar along with the VH, DH and JH segments reported in the sequencing datasets, were synthesized (Genescript) and cloned into pPE2 expression vector by restriction digestion.

#### High-throughput antibody profiling in unselected patient cohort

Small volumes (< 100 µL) of plasma samples were obtained from the Institute of Clinical Chemistry at the University Hospital of Zurich as unique biospecimens. In order to test the samples for the presence of IgGs reactive against human recPrP_23-230_, high-binding 1536-well plates (Perkin Elmer, Spectra Plate 1536 HB) were coated with 1 µg/mL human recPrP_23-230_ in PBS at 37 °C for 1 h, followed by 3 washes with PBS-T and by blocking with 5% milk in PBS-T for 1.5 h. Three µL plasma, diluted in 57 µL sample buffer (1% milk in PBS-T), were dispensed at various volumes into human recPrP_23-230_ coated 1536-well plates using contactless dispensing with an ECHO 555 Acoustic Dispenser (Labcyte). Thereby, dilution curves ranging from plasma dilutions 1:50 to 1:6400 were generated (eight dilution points per patient plasma sample). After the sample incubation for 2 h at RT, the wells were washed five times with wash buffer and the presence of IgGs bound to the human recPrP_23-230_ was detected using an HRP-linked anti-human IgG antibody (Peroxidase AffiniPure Goat Anti-Human IgG, Fcγ Fragment Specific, Jackson, 109-035-098, at 1:4000 dilution in sample buffer). The incubation of the secondary antibody for one hour at RT was followed by three washes with PBS-T, the addition of TMB, an incubation of three minutes at RT, and the addition of 0.5 M H_2_SO_4_. The well volume for each step reached a final of three µL. The plates were centrifuged after all dispensing steps, except for the addition of TMB. The absorbance at 450 nm was measured in a plate reader (Perkin Elmer, EnVision) and the inflection points of the sigmoidal binding curves were determined using a custom designed fitting algorithm. Samples reaching half-maximum saturation (shown as the inflection point of the logistic regression curve) at a concentration ≤ 1:100, i.e. at -log(EC50) ≥ 2, and with a mean squared residual error < 20% of the actual –log(EC50) were considered hits. The inclusion of a threshold for fitting error ensures a reliable identification of positives from high-throughput screening. For the validation screen, hits from HTS were tested against a panel of antigens consisting of human recPrP_23-230_, bovine serum albumin (BSA, Thermo Scientific), and human Apolipoprotein E ε3 (Peprotech), at 1 ug/mL for all antigens. An additional uncoated condition was included to account for residual binding to either the blocking buffer or to the plate. The validation screen was performed identically to the HTS, except that triplicates were used instead of unicates. Samples from the validation screen were considered confirmed if -log(EC50) for PrP ≥ 2 (distinct reactivity against PrP) and -log(EC50) for other targets < 2 (no distinct reactivity against any other control target). All data, including the patient-associated demographic and medical data, was stored in a MS-SQL database. Python and R software as well as GraphPad Prism were used for data visualization and statistical testing. Categorical data was tested with chi-square statistics and Bonferroni correction for multiple comparisons was applied. P-values lower than 0.01 were considered significant.

#### Kappa and Lambda light chain ELISA for total IgG and anti-PrP autoantibodies from plasma

To test total IgG from plasma, 384 well HB plates (Perkin Elmer) were coated with AffiniPure Goat anti-human IgG, Fc gamma fragment antibody, at 1 µg/mL in PBS for 2 h at 37 °C. To test anti-PrP autoantibodies, plates were coated with human recPrP_23-230_. After washing and blocking in 5 % Milk in PBS-T for 90 minutes at RT, serial dilutions of patient plasma (starting from 1:8000 and 1:50 dilution for total IgG and anti-PrP autoantibody ELISA, respectively) were incubated for 2 h at 37°C. Pooled normal human plasma (Innovative Research, #IPLA-N), IgG-Kappa, IgG-Lambda at a concentration of 200 ng/ml or humanized POM1, (hPOM1 at 200 ng/ml and 2 µg/ml for total IgG and anti-PrP autoantibody ELISA, respectively) served as controls. After washing five times, the following HRP-coupled secondary antibodies were incubated at 1:4000 in PBS-T for 1 h, at RT: Goat anti-human Kappa-HRP, goat anti-human Lambda-HRP or goat anti-human IgG Fc-gamma specific-HRP (Jackson ImmunoResearch #109-035-098). The plates were developed as described above.

#### IgG purification from human blood samples

For further validation, 50-250 μl of plasma were used for immunoglobulin purification via 1:1 mixture of proteinA/proteinG resin (GE Healthcare). Slurry was washed once with PBS. Blood samples were spun down at 16’000 x g for 5 min, mixed with 10% of slurry and incubated at 4°C o/n on a rotating wheel. Slurry-blood mix was transferred to screw cap spin columns (Pierce) and beads were washed twice with PBS. Beads were spun down one more time to remove residual PBS and 100 µl of 0.1 M glycine, pH 1.8 was added to the beads. 400 µl of 1 M Tris, pH 8.0 were added to the bottom of the spin columns for neutralization of elution buffer after centrifugation. For buffer exchange, 0.5 ml Centrifugal Filters (Amicon) were used. Protein concentration was measured using NanoDrop ND-1000 Spectrophotometer.

IgG fractions of blood samples were diluted at 100 µg/ml in 1% skim milk in PBS-T and assessed for binding to human recPrP_23-230_, murine recPrP_23-110_ and human recPrP_121-230_ and BSA (coated at 43 nM) in ELISA using goat anti-human IgG Fc-gamma specific-HRP (Jackson ImmunoResearch #109-035-098), 1:4’000 for detection.

## Acknowledgments

We thank Rita Moos and Cinzia Tiberi for support in recombinant antigens and Fab purification, Irina Abakumova for TR-FRET antibody-fluorophore conjugation, and Stefan Schauer for the SPR experiments. We also thank the hospital patients supporting research, especially the individuals who generously agreed to grant us a second blood donation. We acknowledge the help provided by the Clinical Trial Center (CTC), University Hospital of Zurich, and the Institute of Clinical Chemistry, University Hospital of Zurich. AA is the recipient of an Advanced Grant of the European Research Council and grants from the Swiss National Research Foundation, the Nomis Foundation, the Swiss Personalized Health Network (SPHN, 2017DRI17), and a donation from the estate of Dr. Hans Salvisberg. Assunta Senatore and Silvia Sorce are recipient of the Career Development Award grant from the Synapsis foundation.

## Author Contributions

Conceived and designed the experiments: AS NG AA. Performed the experiments: AS KF ME GH JG SF ML RR. Analyzed the data: AS NG AC ME SS. Contributed reagents/materials/analysis tools: NG SE TP SH AA. Wrote the paper: AS SH AA.

---

**Extended data figure S1:**
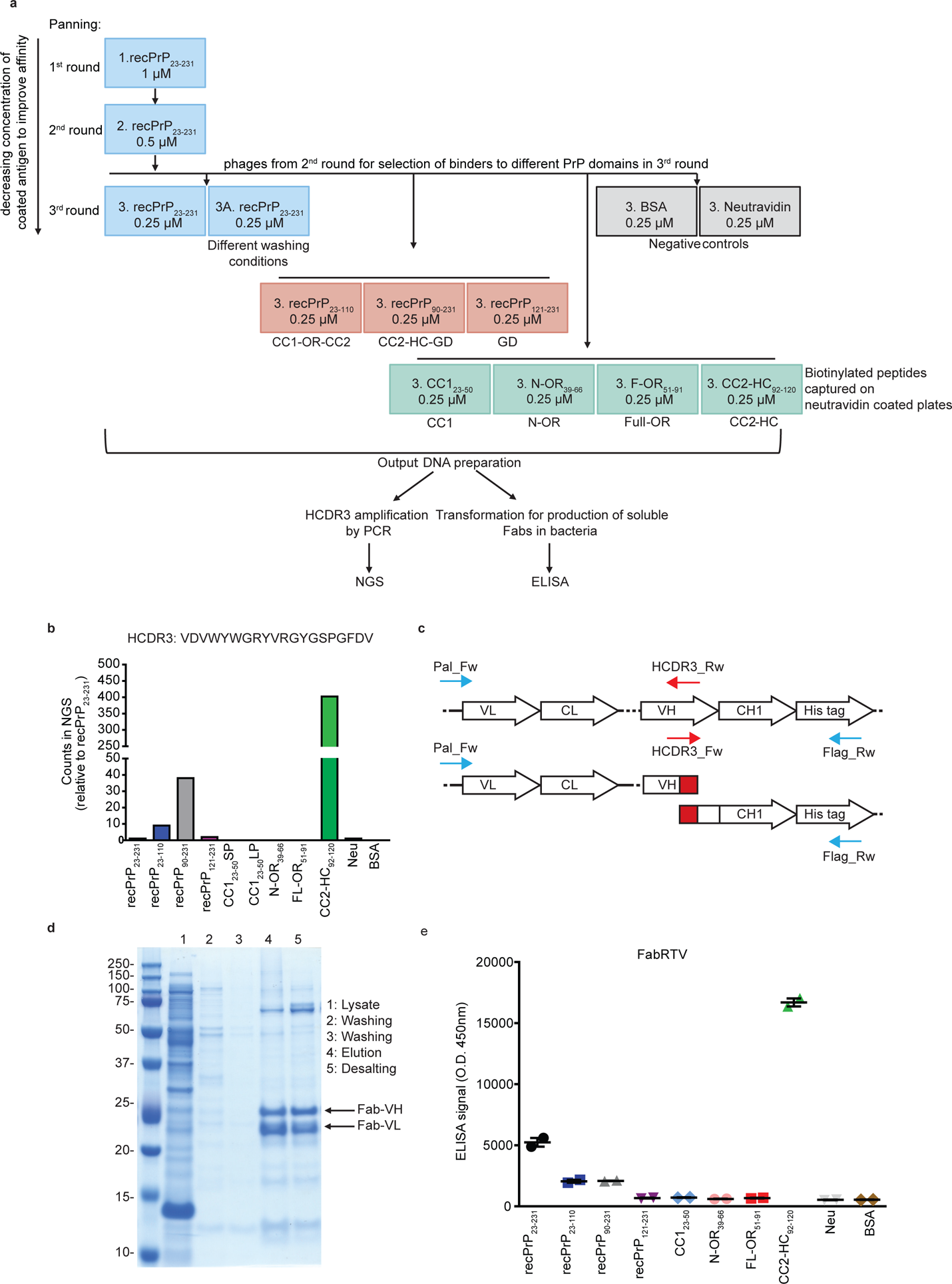
Selection strategy to identify anti-PrP Fabs from a synthetic human Fab phage library and rescue of NGS identified Fab clones. (a) RecPrP_23-231_ (blue boxes) was used as target for the first and second round of panning. In the third round, selected phages were panned against recPrP_23-231_, recPrP_23-231_ under different washing conditions, recPrP fragments (red boxes) and peptides (green boxes) to select for Fabs targeting specific regions of PrP. The DNA preparation from the selected phages was used for NGS or ELISA. Negative controls: BSA and Neu (grey boxes). (b) Bar plot of the NGS counts, relative to recPrP_23-231_, for the indicated HCDR3 sequence. (c) PCR strategy to retrieve the clone of interest. Specific HCDR3_Rw and HCDR3_Fw primers (red arrows) were designed as complementary to the sequence to be rescued. In two separate PCR reactions, the HCDR3_Rw primer was used in combination with the Pal_Fw primer (blue arrow) annealing upstream of the VL, whereas the HCDR3_Fw primer was used in combination with the Flag_RW primer annealing to the His-tag sequence (blue arrow). The two amplicons were then assembled in a second PCR reaction resulting in the full Fab sequence. (d) The retrieved Fab clone (FabRTV) was expressed in *E.coli* and purified by IMAC. The purity of the Fab was analyzed by SDS-PAGE. (e) FabRTV was tested for its binding specificity to recPrP_23-231_ and the indicated PrP fragments and peptides by ELISA. As determined by the NGS analysis (b) and given as an example, FabRTV binds to CC2-HC_92-120_.

**Extended data figure S2:**
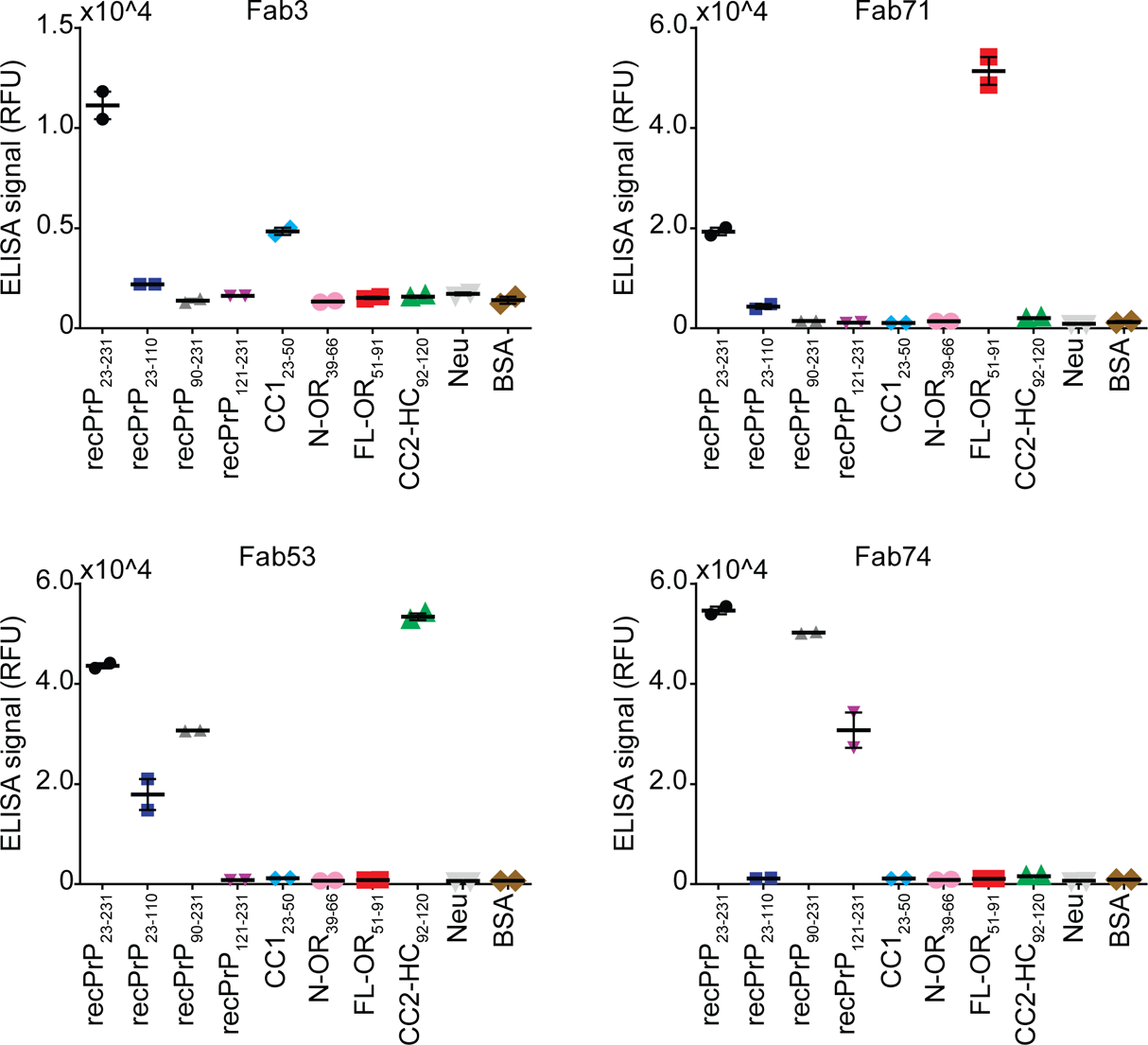
Binding specificity of the retrieved anti-PrP Fabs. ELISA signal (relative fluorescence units, RFU) indicating the binding reactivity of selected, purified anti-PrP Fabs to mouse recPrP_23-231_, mouse recPrP fragments and selected N-terminal PrP peptides. Negative controls: Neu and BSA. Fab3: CC1_23-50_ binder; Fab71: OR_51-91_ binder; Fab53: CC2-HC_92-120_ binder; Fab74: GD binder.

**Extended data figure S3:**
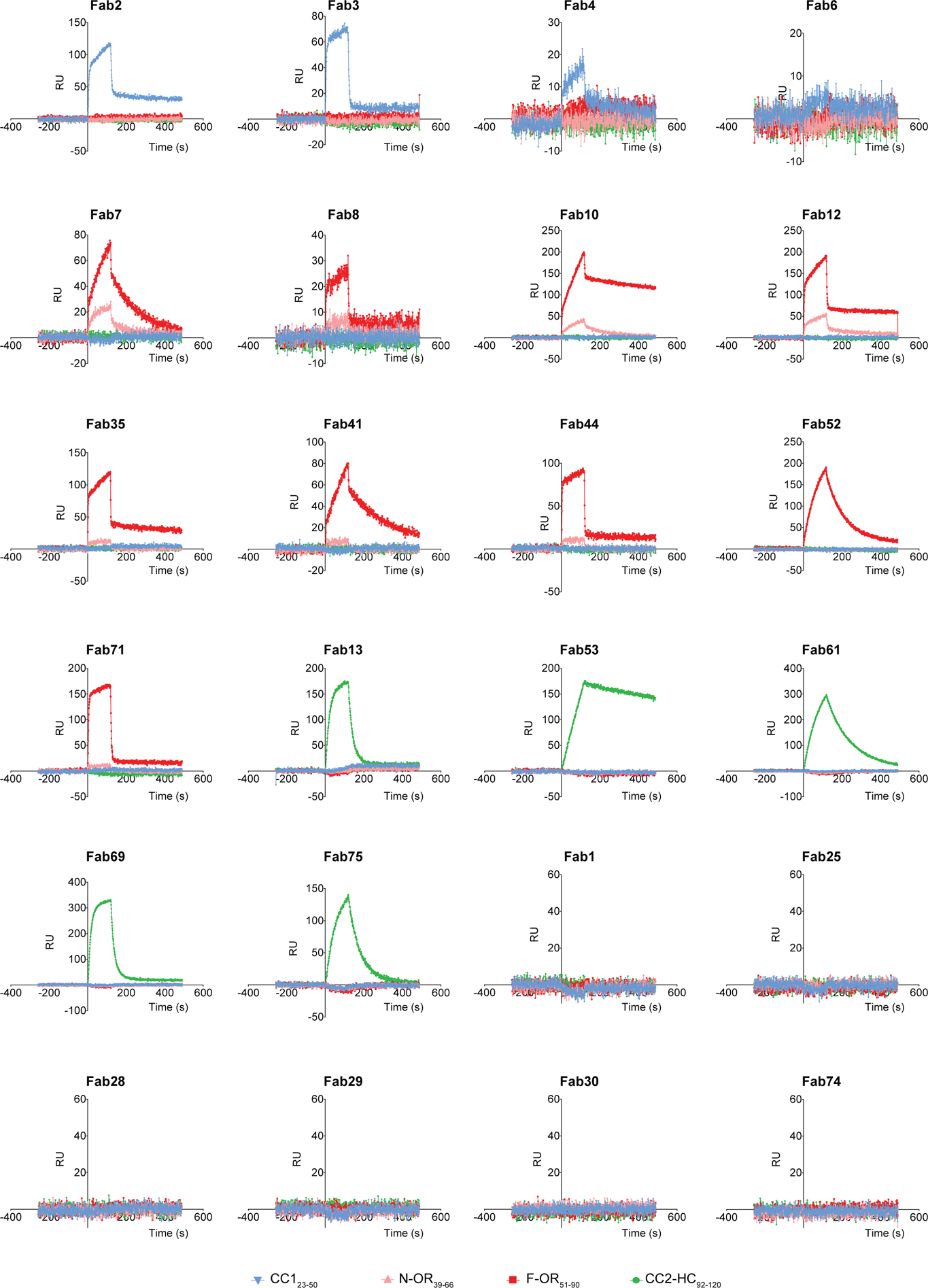
Epitope confirmation by SPR. Biotinylated PrP peptides were immobilized on an NLC sensor chip and the binding of the indicated Fabs was measured by SPR. Epitope assessment by SPR confirmed the results obtained by ELISA. RU: relative units.

**Extended data figure S4:**
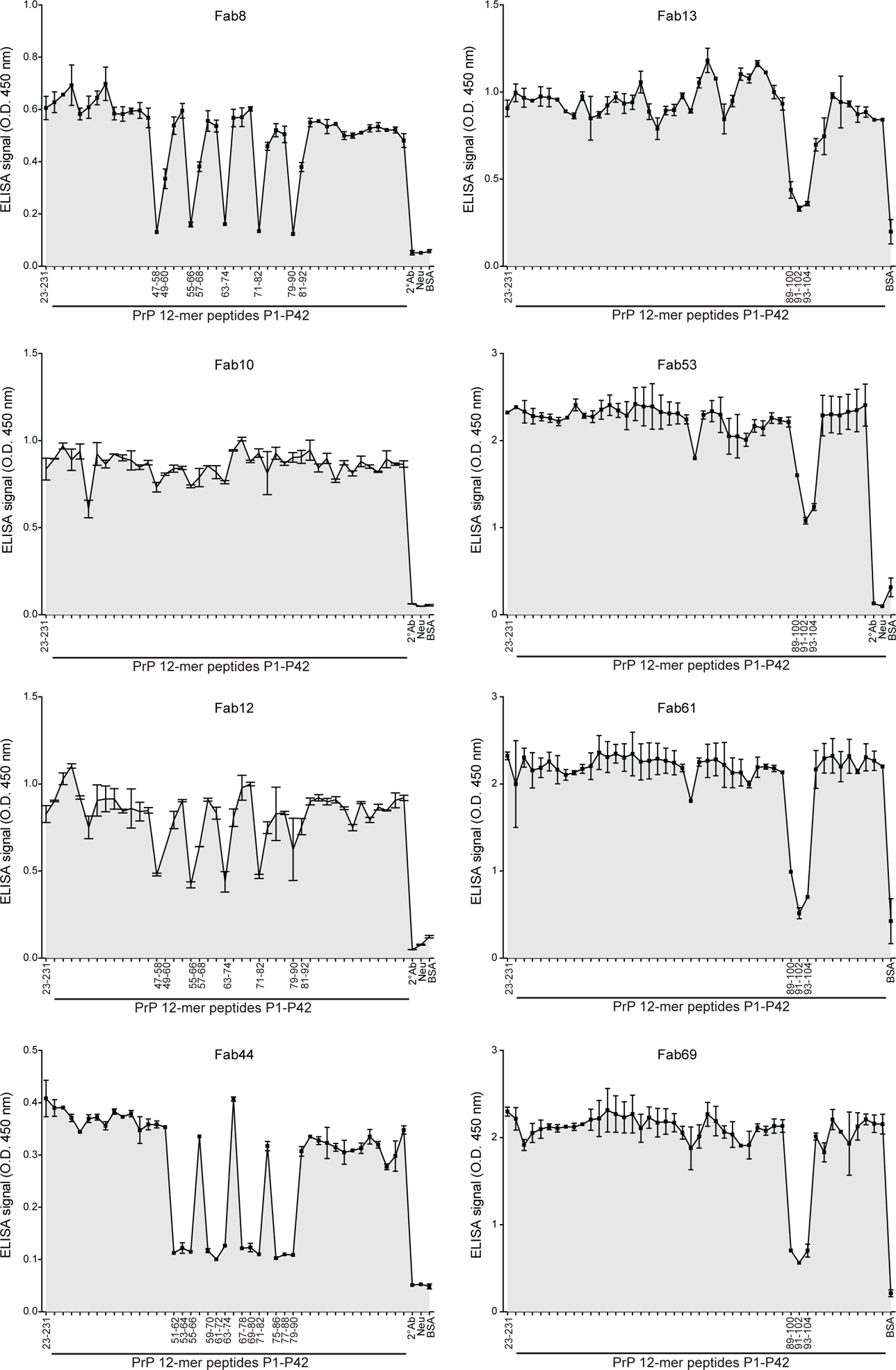
Epitope mapping by competition ELISA. FT-competition ELISA to map the epitopes of the OR_51-91_ binders (Fab8, Fab44, Fab12) and CC2-HC_92-120_ binders (Fab13, Fab61, Fab69). Peptides that strongly inhibit the binding of the Fabs to recPrP_23-231_ are indicated by their residue numbers in the PrP sequence and reflect the respective binding epitopes. Positive control: recPrP_23-231_. Negative controls: Neu, BSA, and 2° Ab.

**Extended data figure S5:**
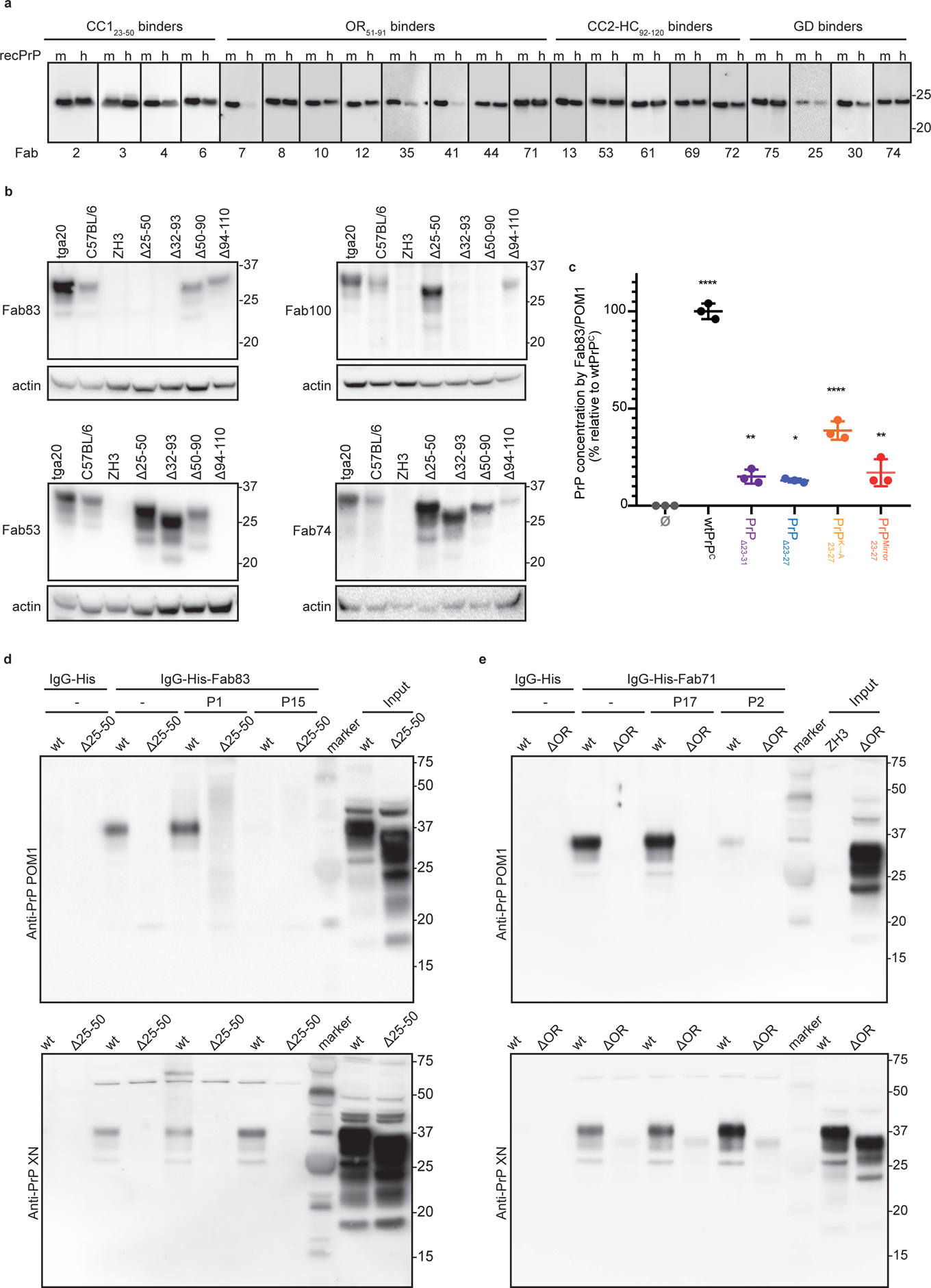
Validation of the reactivity of the anti-PrP Fabs by Western blot, ELISA and immunoprecipitation. (a) Western blot analysis to compare the reactivity of the indicated Fabs to mouse (m) recPrP_23-231_ and human (h) recPrP_23-230_ and **(b)** to full-length and truncated PrP^C^ in BHs of various mouse lines. Actin was used as loading control. **(c)** ELISA to compare the efficiency of Fab83 and POM1 to detect and quantify wt and mutant PrP^C^ levels in CAD5-*Prnp*^-/-^ cells transfected either with wt or mutant PrP^C^ (deletion mutants PrPΔ23-31 or PrPΔ23-27; *PrP*^K→A^_23–27_ with lysine residues 23, 24, and 27 replaced by alanine; *PrP*^Mirror^_23−27_ with KKRPK exchanged to KPRKK). All concentrations (% relative to wtPrP^C^) were determined by interpolating the ELISA signal to a standard curve of mouse recPrP_23-231_. Two-Way ANOVA with Dunnett post-hoc test: * p<0.05; ** p<0.01; *** p<0.001; **** p<0.0001. n = 3 technical replicates. **(d)** Fab83 coupled beads immunoprecipitated wtPrP^C^ from NBH, but not from BH of PrPΔ25-50 mice lacking the respective epitope. WtPrP^C^ specifically eluted by competition with the Fab83 epitope-targeting peptide P1 (residues 23-34), but not with the unrelated peptide P15 (top panel). Elution with a SDS buffer was used as non-specific control (lower panel). Control: IgG-His. Molecular sizes are presented in kDa. **(e)** Same as (d), but for Fab71 coupled beads which immunoprecipitated wtPrP^C^ from NBH, but not from BHs of PrPΔOR mice. WtPrP^C^ specifically eluted with the epitope-targeting peptide P17 (residues 55-66), but not with the unrelated peptide P2 (top panel).

**Extended data figure S6:**
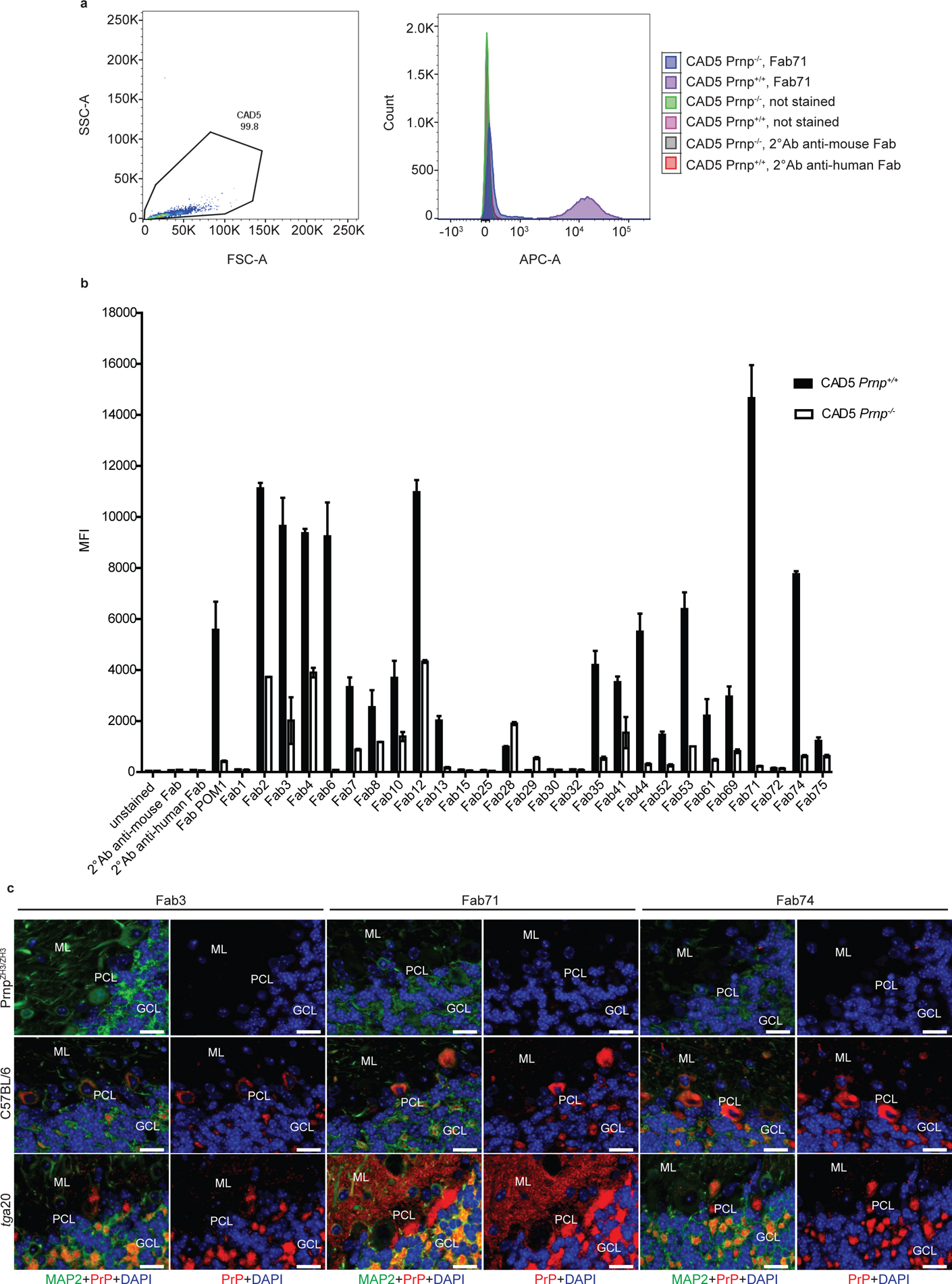
Anti-PrP Fabs detect cell surface exposed wtPrP^C^ and stain wtPrP^C^ in paraffin embedded mouse brain sections. (a) Flow cytometry analysis of the Fabs compared to Fab-POM1 used as positive control for the staining. CAD5 *Prnp^+/+^* and *Prnp^-/-^* cells were incubated with the Fabs at 150 µg/ml, and analyzed with an APC-labelled anti-human Fab antibody. Representative flow cytometry dot plot of FSC-A (forward scatter)/SSC-A (side scatter) of CAD5 *Prnp^+/+^* stained with Fab71 (left) and fluorescence intensity plot of CAD5 *Prnp^+/+^* and *Prnp^-/-^* cells. **(b)** Bar graph showing the average mean fluorescent intensity (MFI) for each Fab. Increased MFIs indicate binding of the Fabs to wtPrP^C^ on the cell surface. Each Fab was tested in duplicate. **(c)** Immunofluorescent staining of cerebellar brain sections of *Prnp*^ZH3/ZH3^, wt and *tg*a20 mice with either Fab3, Fab71, or Fab74 (displayed in red). Anti-MAP2 antibody (microtubule-associated protein 2) was used as neuronal marker (in green) and DAPI to stain the cell nuclei (in blue). The Fabs detected wtPrP^C^ in the cerebellar granule cell layer (CGL) and molecular layer (ML) of wt and *tg*a20 mice. As expected, the Fabs did not detect wtPrP^C^ in Purkinje cell layer (PCL) of *tg*a20 mice. The higher staining intensity of Fab71 might be caused by its ability to recognize the repetitive epitopes within the OR multiple times. Scale bar: 20 µm.

**Extended data figure S7:**
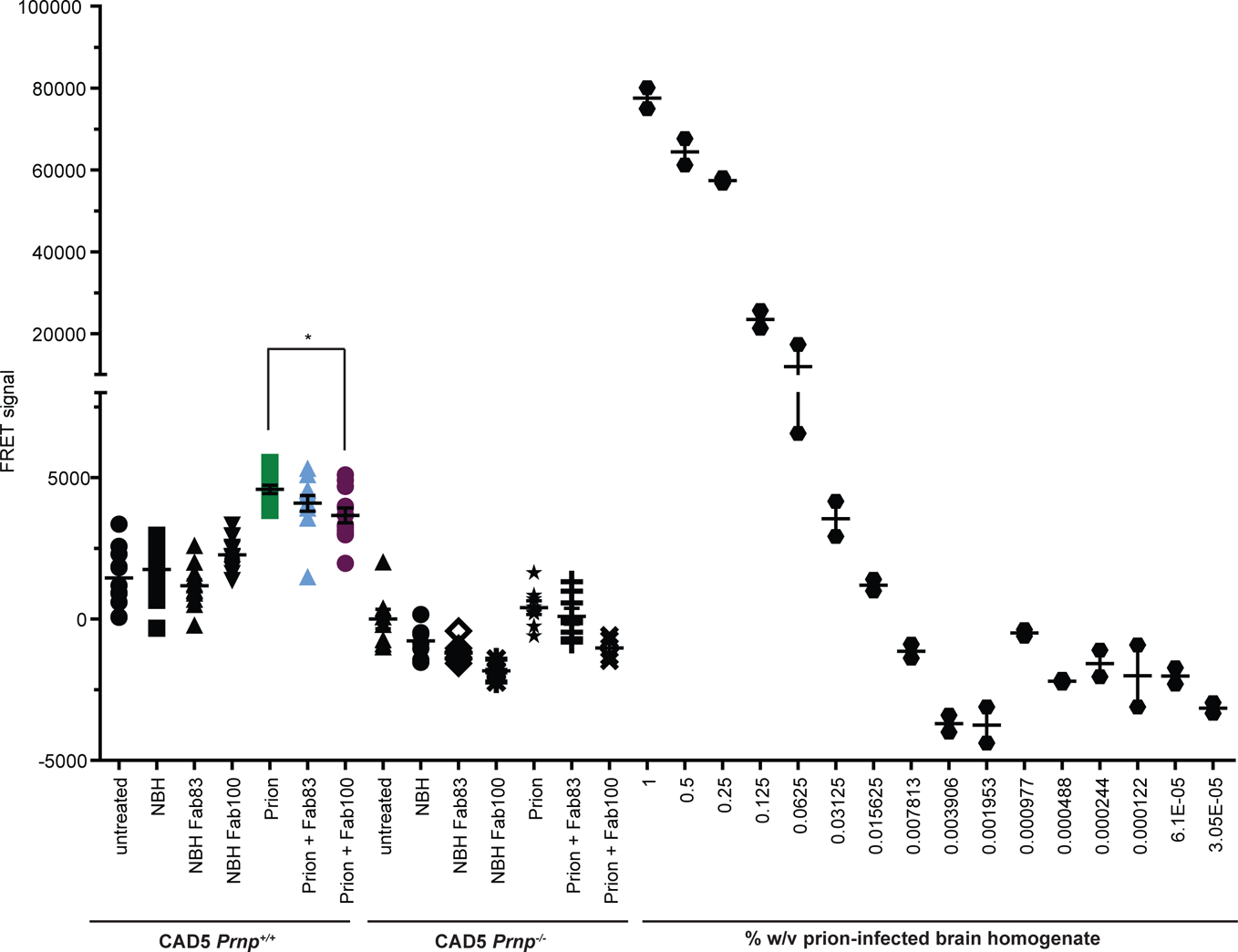
TR-FRET for the detection of wtPrP^Sc^ in prion-infected CAD5 *Prnp^+/+^* after treatment with Fab100 and Fab83. Treatment of prion-infected CAD5 *Prnp^+/+^* cells with Fab100 (OR_51-91_ binder, * p<0.05), but not with Fab83 (CC1_23-50_ binder), reduced wtPrP^Sc^ levels compared to untreated prion-infected CAD5 *Prnp^+/+^* cells. Negative controls: NBH and CAD5 *Prnp^-/-^* cells. The dilution range (3×10^-5^ % to 1% w/v) of prion-infected BH is shown on the right site to indicate the linear range of the assay. The FRET signal of untreated CAD5 *Prnp^-/-^* cells was set to zero. The FRET anti-PrP antibody pair POM19-Eu and POM1-APC was used for detection. For prion infected CAD5 *Prnp^+/+^* cells, the assay was performed in biological triplicates represented as four technical replicates for Fab100 and Fab83. Each dilution of prion-infected BH is represented in technical duplicates. One-way ANOVA, Bonferroni’s multiple comparisons was used for statistical analysis.

**Extended data figure S8:**
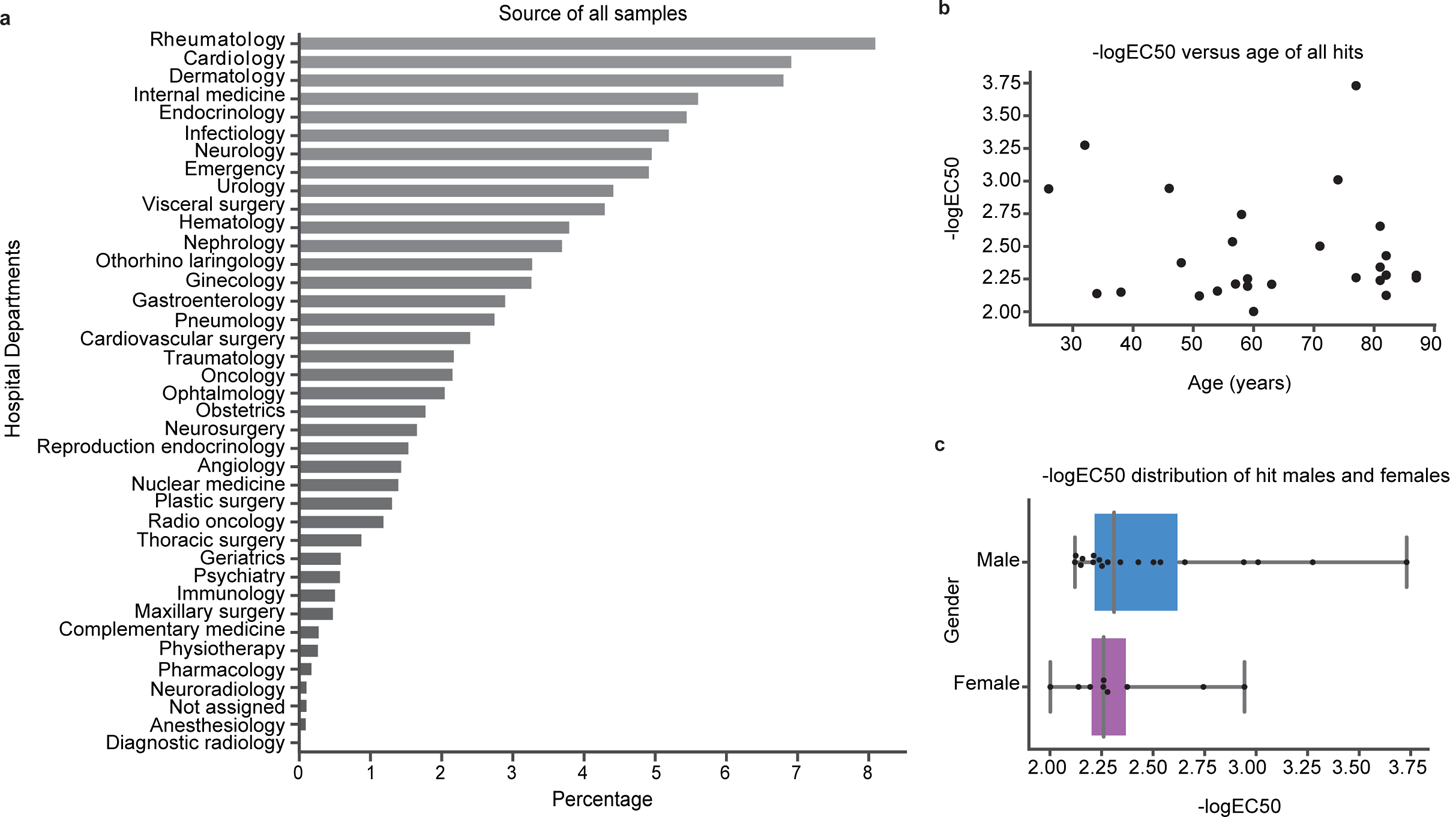
Medical and demographic data of the hospital cohort screened for anti-PrP antibodies and of the hits identified in the primary screening. (a) Bar plot indicating the hospital departments from which the tested 37,894 samples were obtained. (b-c) Correlations between -log(EC50) and age (b), -log(EC50) and sex (c). Black dots in panels (b) and (c) indicate each of the 27 hits that were identified in the primary screen.

**Table S1:** **Matrix of the different washing conditions during panning**

| **Washing time** | **A1 (standard)** | **A2 (high salt)** | **A3 (detergent)** |
| --- | --- | --- | --- |
| 5 times quick,<br>10 times 5 min | PBST | PBST + 1M NaCl | PBS + 2% Octyl β-D-glucopyranoside |
| 5 times quick,<br>7 times 5 min | PBS | PBS + 1M NaCl | PBS + 2% Octyl β-D-glucopyranoside |
| 3 times 5 min | PBS | PBS | PBS |

**Table S2:** Criteria for sorting the NGS counts for epitope prediction of the anti-PrP Fabs

| Panning output NGS libraries |  |  |  |  |  |  |  |  |  |  |  |  |  |  |  |
| --- | --- | --- | --- | --- | --- | --- | --- | --- | --- | --- | --- | --- | --- | --- | --- |
|  |  |  | Ref | binding strength |  | synthetic biotinylated PrP peptides |  |  |  |  | recombinant PrP fragments |  |  | negative controls |  |
| mouse PrP amino acid residues | PrP domains |  | recPrP<sub>23-231</sub> | recPrP<sub>23-231</sub> high salt | recPrP<sub>23-231</sub> detergent | CC1<sub>23-50</sub> SP | CC1<sub>23-50</sub> LP | N-OR<sub>39-66</sub> | F-OR<sub>51-91</sub> | CC2-HC<sub>92-120</sub> | recPrP<sub>23-110</sub> | recPrP<sub>90-231</sub> | recPrP<sub>121-231</sub> | Neutravidin | BSA |
| 23-38 | CC1 | CC1_SP | 1 | ns | ns | >0 | ns | 0 | 0 | 0 | >0 | 0 | 0 | 0 | 0 |
|  |  | CC1_LP | 1 | ns | ns | ns | >0 | 0 | 0 | 0 | >0 | 0 | 0 | 0 | 0 |
| 39-50 | CC1-N-OR |  | 1 | ns | ns | ns | ns | >0 | 0 | 0 | >0 | 0 | 0 | 0 | 0 |
| 50-66 | N-OR |  | 1 | ns | ns | 0 | 0 | >0 | ns | 0 | >0 | 0 | 0 | 0 | 0 |
| 51-91 | F-OR |  | 1 | ns | ns | 0 | 0 | ns | >0 | 0 | >0 | 0 | 0 | 0 | 0 |
| 92-110 | CC2-HC |  | 1 | ns | ns | 0 | 0 | 0 | 0 | >0 | >0 | >0 | 0 | 0 | 0 |
| 111-120 | CC2-HC |  | 1 | ns | ns | 0 | 0 | 0 | 0 | >0 | 0 | >0 | 0 | 0 | 0 |
| 90-231 | CC2-GD |  | 1 | ns | ns | 0 | 0 | 0 | 0 | 0 | 0 | >0 | >0 | 0 | 0 |
| 121-231 | GD |  | 1 | ns | ns | 0 | 0 | 0 | 0 | 0 | 0 | 0 | >0 | 0 | 0 |

Ref: reference panning to recPrP<sub>23-231</sub>
CC1\_SP: CC1 Solid Phase
CC1\_LP: CC1 Liquid Phase
ns: not selected

**Table S4:**
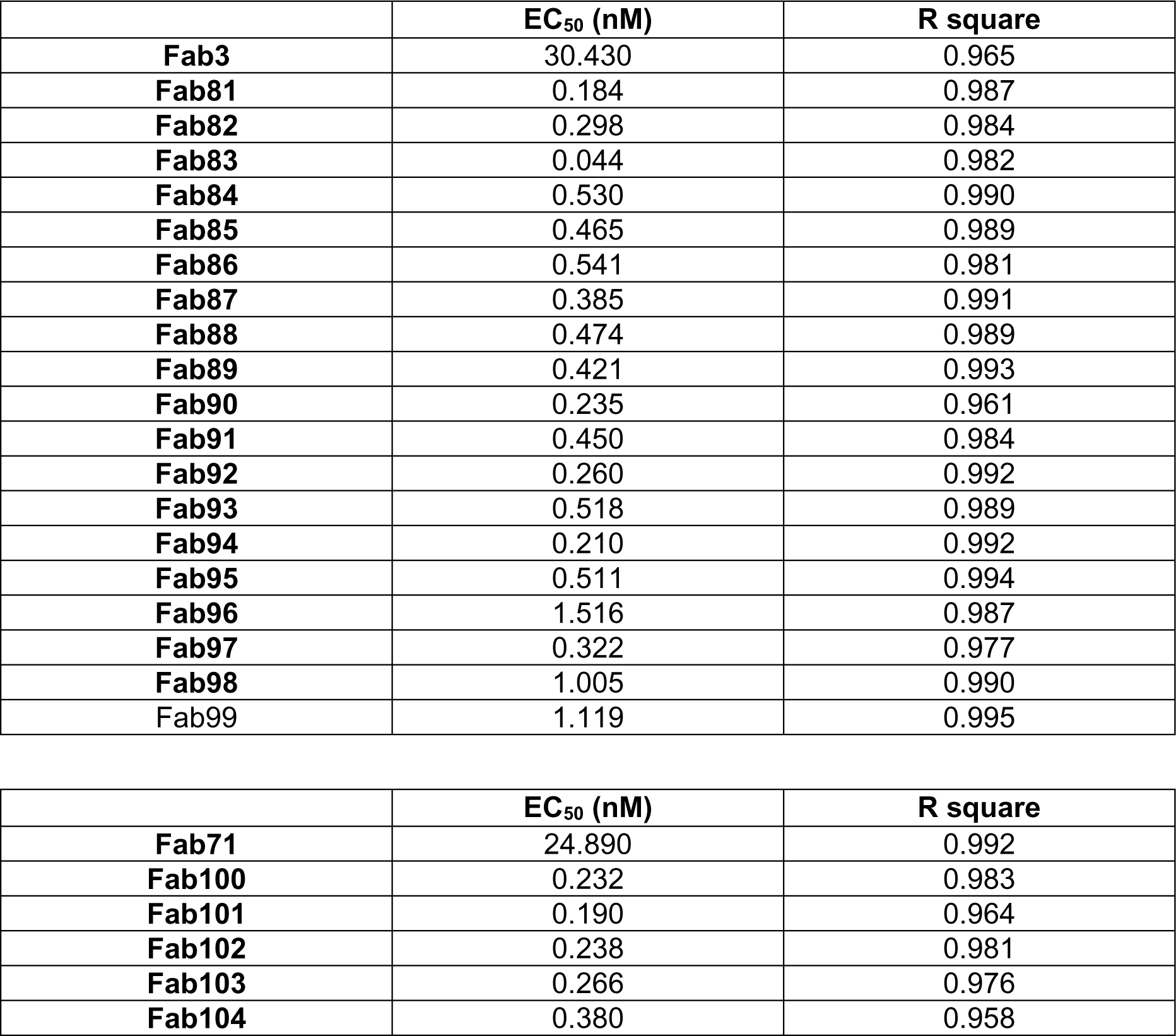
**EC50 of Fab3 and Fab71 and their respective affinity matured variants as determined by ELISA**

|  | **EC<sub>50</sub> (nM)** | **R square** |
| --- | --- | --- |
| **Fab3** | 30.430 | 0.965 |
| **Fab81** | 0.184 | 0.987 |
| **Fab82** | 0.298 | 0.984 |
| **Fab83** | 0.044 | 0.982 |
| **Fab84** | 0.530 | 0.990 |
| **Fab85** | 0.465 | 0.989 |
| **Fab86** | 0.541 | 0.981 |
| **Fab87** | 0.385 | 0.991 |
| **Fab88** | 0.474 | 0.989 |
| **Fab89** | 0.421 | 0.993 |
| **Fab90** | 0.235 | 0.961 |
| **Fab91** | 0.450 | 0.984 |
| **Fab92** | 0.260 | 0.992 |
| **Fab93** | 0.518 | 0.989 |
| **Fab94** | 0.210 | 0.992 |
| **Fab95** | 0.511 | 0.994 |
| **Fab96** | 1.516 | 0.987 |
| **Fab97** | 0.322 | 0.977 |
| **Fab98** | 1.005 | 0.990 |
| **Fab99** | 1.119 | 0.995 |
| **Fab71** | 24.890 | 0.992 |
| **Fab100** | 0.232 | 0.983 |
| **Fab101** | 0.190 | 0.964 |
| **Fab102** | 0.238 | 0.981 |
| **Fab103** | 0.266 | 0.976 |
| **Fab104** | 0.380 | 0.958 |

**Table S5:** Features and clinical data of the subjects displaying high titre of anti-PrP autoantibodies in the plasma.

| Patient ID | Core diagnoses | Sex | Year of birth |
| --- | --- | --- | --- |
| Patient_01 | Valvular and coronary heart disease; chronic obstructive pulmonary disease | Male | 1943 |
| Patient_02 | Contusion of digit | Female | 1971 |
| Patient_03 | Chronic lymphatic leukaemia; valvular cardiac disease; chronic kidney insufficiency; oncocyoma | Male | 1937 |
| Patient_04 | Not shown |  |  |
| Patient_05 | Myeloid leukaemia | Male | 1959 |
| Patient_06 | Aortic stenosis | Male | 1947 |
| Patient_07 | Not shown |  |  |
| Patient_08 | Aortic aneurysm and dissection | Male | 1937 |
| Patient_09 | Ulcerating colitis; drug abuse | Female | 1970 |
| Patient_010 | Not shown |  |  |
| Patient_014 | HIV infection; erectile dysfunction; gonarthrosis | Male | 1959 |
| Patient_015 | Intraspinal space-consuming lesion (ganglioneurom WHO grade I); drug abuse | Male | 1985 |
| Patient_016 | Coronary heart disease; severe kidney insufficiency; hypothyreosis | Female | 1959 |
| Patient_017 | Polyserositis; mamma carcinoma | Female | 1958 |
| Patient_018 | HIV infection | Male | 1991 |
| Patient_019 | Not shown |  |  |
| Patient_020 | Urothel carcinoma; adenocarcinoma; arrhythmogenic cardiac disease | Male | 1936 |
| Patient_021 | Coronary 2-vessel disease | Male | 1954 |
| Patient_022 | Coronary heart disease; neuropathic pain | Male | 1935 |
| Patient_023 | T cell lymphoma | Male | 1967 |
| Patient_024 | Injury of the pelvic ring | Male | 1964 |
| Patient_025 | Status epilepticus; myxofibrosarcoma; slight dementia type Alzheimer's disease | Female | 1931 |
| Patient_026 | Squamos cell carcinoma | Female | 1941 |
| Patient_027 | Not shown |  |  |
| Patient_028 | Syncope; severe depressive episodes | Female | 1985 |
| Patient_029 | Not shown |  |  |
| Patient_030 | Adenocarcinoma; signs of dementia | Male | 1937 |

Shown are the core diagnoses, the sex, and the year of birth for the 21 out of 27 individuals identified with autoantibodies against recombinant PrP in the high-throughput antibody profiling. We have refrained from displaying any patient-associated information from patients who do not gave expressive consent by signing the hospital-wide general consent of the University Hospital of Zurich. While many but not all of the patients included here are highly multimorbid, only the core diagnoses were selected for display, also with respect to the appropriate handling of sensitive patient data.

**Table S6:** **Comparison of age, gender and clinical data between subjects with and without high titre PrP autoantibodies**.

| Median age hits | # hits | Median age non-hits | # non-hits | P-value |
| --- | --- | --- | --- | --- |
| 61 | 27 | 55 | 48691 | 3.58E-02 |

| % male hits | % female hits | % male non-hits | % female non-hits | P-value |
| --- | --- | --- | --- | --- |
| 66.67 | 33.33 | 52.42 | 47.64 | 1.96E-01 |

| ICD-10 code | # hits | # non-hits | # hits | # non-hits | P-value | Bonferroni |
| --- | --- | --- | --- | --- | --- | --- |
| E87 | 6 | 4830 | 22.22 | 9.92 | 4.56E-02 | 1.82E-01 |
| I25 | 7 | 6207 | 25.93 | 12.75 | 7.33E-02 | 2.93E-01 |
| I10 | 9 | 11595 | 33.33 | 23.81 | 2.59E-01 | 1 |
| N18 | 5 | 5874 | 18.52 | 12.06 | 3.76E-01 | 1 |

Statistical testing for difference in age (median test, significance at $\alpha \leq 0.01$ ) or gender (chi-square test, significance at $\alpha \leq 0.01$ ) revealed no significant difference. Analysis of enriched ICD-10 codes in patients with anti-PrP-autoantibodies versus patients without. No patterns could be discerned and no ICD-10 code reached a statistically significant enrichment (chi-square test, significance at $\alpha \leq 0.01$ , Bonferroni correction for multiple comparisons).

